# Slow diffusion limits phosphorylation in a biomolecular condensate

**DOI:** 10.64898/2026.01.29.702494

**Authors:** Nicolas S. Gonzalez-Foutel, Ankush Garg, Evi Setiani Lande, Assia Khalild, Line Mørkholt Lund, Victoria Birkedal, Chloé Martens, Magnus Kjaergaard

## Abstract

Biomolecular condensates form dynamic compartments that regulate biochemical reactions in cells. Condensates recruit many kinases and regulate their enzymatic activity. Condensates alter the rate of enzymatic reactions through several opposing effects, so it is unclear whether these mostly enhance or retard phosphorylation. Here, we use a synthetic condensate formed by intrinsically disordered proteins to show that slow diffusion in the condensate controls phosphorylation kinetics in the dense phase. We vary the length of substrates by appending phase-separating repeat proteins of different lengths, in order to study how phosphorylation depends on partitioning, diffusion and volume fraction across substrate motifs with different intrinsic kinetics. The condensate environment is generally inhibitory to phosphorylation, although the enzyme remains intact. This inhibition is partially offset by an enhanced reaction rate in the dilute phase, likely due to soluble nanoclusters. Phosphorylation rates are strongly correlated to diffusion coefficients of substrates in the condensate, suggesting mass-transport limitation. Our results suggest that condensates can modify the substrate usage of a kinase via different trade-offs between diffusion and partitioning. We suggest that diffusion limitations are likely a common feature of many macromolecular reactions in condensates, and that high fluidity is crucial for condensates to act as reaction crucibles.

## Introduction

Condensates are formed by phase separation into a dense phase - with a high concentration of the scaffold macromolecule - and a dilute phase with a low concentration.^1^ Condensates can permit entrance of other molecules – clients – and either increase or decrease their concentrations depending on interactions between the scaffold and clients. A picture of the interior of condensates is emerging, where their architecture mirrors the complexity and diversity of macromolecular constituents. Formation of biomolecular condensate can act as biochemical switches by increasing or reducing the rates of biochemical reactions. Biochemical reactions in condensates are affected by many factors, including partitioning of enzymes and substrates, an effective solvent environment with different pH^2,3^ and polarity,^4,5^ and changes in the structure and dynamics of macromolecules.^6,7^ These factors may affect an enzyme in opposing directions and are likely to differ between condensates and enzymes.^8^

Phosphorylation has a bidirectional effect on condensates: Phosphorylation of condensate scaffolds alters their physicochemical properties and interactions, and in turn their propensity to phase separate.^9,10^ Phosphorylation can trigger both formation or dissolution of condensates. Conversely, many kinases form condensates or enter condensates as clients.^11^ Machine learning algorithms predict that between 13-44% of human kinases have a strong tendency to undergo phase separation.^12^ For example, the regulatory subunit of protein kinase A (PKA) forms condensates in response to cAMP, which concentrates catalytic activity into foci.^13^ Abolition of this phase separation in fusion oncoprotein leads to increased cell proliferation. In contrast, several other fusion oncoproteins form condensates leading to aberrant phosphorylation.^14–16^

Synthetic biology has proven a successful strategy for interrogating the effects of condensates on enzymes. By reconstituting condensates and embedding enzymes therein, enzymatic rates can be studied in vitro and in model cells in response to perturbations of the condensate. Model condensates are typically either formed by multivalent protein-interaction domains^17^ or intrinsically disordered regions,^18,19^ which can drive robust phase separation in vitro and in cells. The enzyme can either be targeted to the condensate through fusion to the condensate scaffold^5^ or recruited as a sub-stoichiometric client. The latter approach results in a non-negligible enzymatic activity in the dilute phase, which has to be accounted for separately.^20^

Recruitment of a MAP kinase to a synthetic condensate led to enhanced phosphorylation in vitro and in cells.^21^ The enhancement was greatest for substrates without a cognate docking site, leading to a relaxation of substrate specificity and to hyper-phosphorylation indicative of processivity.^21^ The rate enhancement was also observed for phosphorylation of tau protein when induced to phase separate by crowding agents.^21^ Likewise, homotypic phase separation accelerated trans-autophosphorylation in tyrosine kinases, as evidenced by a sharp increase in phosphorylation above the saturation concentration.^12^ Molecular dynamics simulations suggest that phosphorylation of TDP-43 by CK1δ occurs in the dense phase, with a largely similar substrate specificity determined mainly by the contact frequency within the condensate.^22^

Here, we study phosphorylation of protein kinase A (PKA) recruited as clients to a condensate consisting of resilin-like repeat proteins.^18^ In contrast to similar studies, we focus on altering the properties of the substrate by embedding PKA substrate motifs into repeat proteins of different length and measuring phosphorylation rates as a function of volume fraction of condensate. The condensates recruit substrate and enzyme, and the enzyme remains structurally intact. In contradiction to predictions based on mass action, we find that the condensates generally retard phosphorylation. Dense phase phosphorylation rates are strongly correlated to diffusion constants of the substrate, which suggest the presence of mass transport limitations. Condensates affect the same substrate motif differently depending on which protein context it is embedded into, which provides a mechanism for differential regulation of substrate usage by kinase. As diffusion and partitioning of clients are both governed by interactions with the condensate scaffold,^23^ this suggests an inherent feature of enzyme reactions in condensates.

## Materials and Methods

### DNA and peptide synthesis

All the plasmids were *de novo* synthesized and codon optimized for expression in *E. coli* (Genscript). The genes of interest were cloned into pET15b vectors between NdeI and BamHI restriction sites. The sequence of all peptides and proteins used are listed in **Supplementary Table 1**. All short peptides were synthesized by Fmoc chemistry at >99% purity (Genscript) and later resuspended in ultrapure water and quantified by absorbance at 280 nm. These peptides are listed in **Supplementary Table 1**.

### Protein expression and purification

The catalytic domain of PKA linked to MBD2 coiled-coil domain (MBD2-PKAc) and His-tag at the N-terminus was expressed in C41(DE3) cells in Luria-Bertani Broth supplemented with 100 μg/mL ampicillin at 37 °C and shaking at 100 rpm as described previously.24 6 x 1L cultures were induced with 1 mM IPTG at OD_600_= 1 and the temperature changed to 20 °C for overnight expression. Cells were harvested by centrifugation (10 minutes at 4000g). The pellet was resuspended in buffer 20 mM NaH_2_PO_4_, 0.5 M NaCl, 5 mM imidazole, 0.1 mM TCEP, 0.2 mM PMSF, 50 mg/L chymostatin, 50 mg/mL leupeptin and 50 mg/L pepstatin at pH 7.5, and lysed by sonication (50% cycle, 70% power for 5 minutes). The lysates were centrifuged for 20 minutes at 14.000 × g and the supernatant was loaded onto a gravity flow column with Ni-NTA superflow (QIAGEN). The column was previously equilibrated with buffer 20 mM NaH_2_PO_4_, 0.5 M NaCl, 0.1 mM TCEP, 5 mM imidazole at pH 7.5 and, once loaded with the supernatant, it was washed with the same buffer containing 20 mM imidazole. The fraction containing the protein of interest was eluted with the same buffer containing 500 mM imidazole. This protocol was enough to obtain MBD2-PKAc with purity above 90% (**Supplementary Fig 1**), so the elution fraction was dialyzed against buffer 20 mM tris base, 150 mM NaCl, 0.1 mM TCEP at pH 7.5 and concentrated to final concentration (150 µM), aliquoted, and stored at -20 °C.

The scaffold proteins ([Q_5,8_]-20 and [Q_5,8_]-20-p66α) and substrates ([Q_5,8_]-x-S_WT, R-3K_) imbedded into multiple resilin-like peptide regions were expressed in BL21(DE3) cells in Terrific Broth supplemented with 100 μg/mL ampicillin at 37 °C and shaken at 100 rpm.^25^ 6 x 1L cultures (per construct) were inoculated with 10mL of saturated preculture and after 4 hours of growth (OD_600_ > 2) induced with 0.8 mM IPTG at the same temperature (37 °C) for overnight expression. Cells were harvested by centrifugation (10 minutes at 4000 ×g). The pellets were resuspended in 30 mL of MQ water per liter of culture harvested and heat-lysed at 90 °C for 1 hour. The lysates were immediately incubated on ice for 30 minutes to cool down and allow the formation of scaffold protein condensates at low temperatures. Then the lysates were centrifuged for 30 minutes at 20.000 g. The soluble fractions were discarded and the insoluble fractions were recovered and resuspended in Binding buffer 20 mM tris base, 0.5 M NaCl, 6 M urea, 5 mM imidazole, pH 7.5 and stirred overnight. The new resuspensions were centrifuged again for 30 minutes at 20.000 ×g and the supernatants recovered. The supernatants were filtered with Whatman paper (Cytiva) and loaded onto a gravity flow column with Ni-NTA superflow (QIAGEN). The columns were previously equilibrated with buffer 20 mM tris base, 0.5 M NaCl, 6 M urea, 5 mM imidazole at pH 7.5 and, once loaded with the supernatant, they were washed with the same buffer containing 20 mM imidazole. The fractions containing the proteins of interest were eluted with the same buffer containing 500 mM imidazole. The eluates were dialyzed against MQ water and the formation of condensates inside the dialysis bags were observed. After 3 days of dialysis and multiple changes of MQ water the condensates formed a film of protein in the walls of the dialysis bags, which were scratched and recovered. The sticky material of pure protein obtained for each construct was then lyophilized and preserved as a powder until use. Proteins presented purity above 90%, assessed by SDS-PAGE (**Supplementary Fig. 1**).

### Temperature dependent turbidity measurements

1ml of [Q_5,8_]-20 at 9ομM was doped with [Q_5,8_]-20-p66α 10 μM (molar ratio of 9:1). The effect of [Q_5,8_]-20-p66α doping on the phase behavior of [Q_5,8_]-20 was evaluated using temperature-dependent turbidity measurements. To investigate the influence of substrate on the phase behavior of the [Q_5,8_]-20 scaffold ([Q_5,8_]-20+[Q_5,8_]-20-p66α scaffold mixture) at 100 μM, [Q_5,8_]-x-S_WT_ (x: 10, 24 and 40) were added at a concentration of 2 μM, and turbidity measurements were performed in their presence and absence (**Supplementary Fig. 2**).

Temperature-dependent turbidity experiments were conducted using a Labbot UV– visible spectrophotometer (Probation Labs, Sweden AB). Optical density at 600 nm (OD_600_) was monitored during programmed cooling from 70 °C to 10 °C in 5 °C increments. This approach exploits the upper critical solution temperature (UCST) behavior of [Q_5,8_]-20, which remains homogeneous at elevated temperatures but undergoes association and phase separation upon cooling, resulting in increased turbidity.

### Phosphorylation Phos-Tag gels

Phosphorylation of [Q_5,8_]-24 S_WT_ was analyzed using SuperSep Phos-tag gels (50 μmol/L) (Fujifilm Wako) 12.5%. The reaction was performed by adding 1.4 µL of 100 mM ATP (final concentration 1 mM) to 140 μL of reaction volume containing 4 μM [Q_5,8_]-24 S_WT_ (0.1 μg/μL), 50 μM [Q_5,8_]-20 and 20 nM MBD2-PKAc in buffer 10 mM Tris:Base, 150 mM NaCl, 10 mM MgCl_2_, 1 mg/ml BSA pH 7.5 (no EGTA) after 30 minutes incubation at 30 °C. Fractions of 20 µL of reaction mix were taken right before adding ATP and after 2, 3, 4, 5, 7, 9 minutes, mixed with 10 µL Phos-tag loading buffer and flash frozen. The control reaction was performed similarly, but without [Q_5,8_]-20. Samples were loaded in SuperSep Phos-tag gel, run for 3.5 hours and revealed with Coomassie Blue staining (**Supplementary Fig. 3**).

### Sample preparation and deuterium labeling

Recruitment of MBD2-PKAc into [Q_5,8_]-20 condensates was performed at 200 µM of [Q_5,8_]-20, 20 µM of [Q_5,8_]-20-p66α and 20 µM of MBD2-PKAc in TBS (20 mM tris-HCl, 150 mM NaCl pH 7.6) for 5 minutes. After the recruitment process, samples were labelled at 20ºC by adding 200 µL of deuterium buffer (20 mM Tris-HCl, 150 mM NaCl pD 7.6) into 150 µL of condensates and enzyme mixture. Labeling reaction was quenched at 30, 90, 300, and 900 seconds by taking 70 µL from the mixture and adding 70 µL of ice cold-quenching buffer (1.5 M guanidine-HCl, 50 mM K_2_HPO_4_, 5 mM KH_2_PO_4_, 0.1% v/v formic acid pH 2.2). The quenched mixture was directly flash frozen in liquid nitrogen and stored at –80ºC. A solution containing 20 µM of MBD2-PKAc in TBS without condensate scaffolds was used as a reference.

### HDX-MS analysis

Thawed samples were directly injected into Waters NanoAcquity UPLC installed with a digestion column of Enzymate BEH Pepsin (Waters) at 200 µL.min^-1^ and 20ºC. Upon digestion, protein samples were trapped for 3 minutes on an Acquity UPLC BEH C18 VanGuard Pre-column (Waters). The RP chromatography was performed on Acquity UPLC BEH C18 (Waters) with linear gradient elution (3-45% gradient of 0.1% formic acid in acetonitrile) at 40 µL.min^-1^. Identification of peptides including the deuterated ones were conducted on SynaptG2 using positive electrospray ionization, independent acquisition, and triwave ion-mobility for improved resolution and identification. Sodium formate was used as a calibration agent and leucine enkephalin for mass accuracy. The pepsin column was washed 5-6 times after injection using pepsin wash buffer (1.5 M guanidinium HCl, 4% v/v acetonitrile, 0.8% v/v formic acid) and a cleaning run was performed after every sample run to avoid clogging the column. Deuterium uptake of all samples was analyzed using DynamX software from PLGS output. The statistical significance of uptake difference between PKA with and without condensates was then checked in Deuteros 2.0.^26^

### Steady-state kinetics

Steady-state experiments to obtain the phosphorylation reaction rates of the homogeneous system were performed by mixing 0.2 nM MBD2-PKAc and 2 μM [Q_5,8_]-X-Substrate in reaction buffer 50 mM tris base, 0.1 mM EGTA, 10 mM magnesium acetate, 150 mM NaCl, pH 7.6. To obtain the total reaction rate of the heterogenous system, we also added to the mixture [Q_5,8_]-20 scaffold at increasing concentrations (4.5, 9, 22.5, 45 and 90 μM) and the tenth part of [Q_5,8_]-20-p66α scaffold (0.5, 1, 2.5, 5 and 10 μM) (9:1), to obtain total scaffold concentrations of 5, 10, 25, 50, and 100 μM, respectively. To obtain the reaction rate corresponding to the dilute phase of the heterogenous system, we followed the same procedure as described above, but after the mixing the samples were ultracentrifuged at 135.000 × g at 20 °C for 30 min and then the supernatant recovered. For all cases, we manually added [γ-^32^P]-ATP (final concentration 0.1 mM of 100-200 c.p.m. pmol^-1^) to a final volume of 200 μL and incubated at 30°C. The moment of ATP addition was considered the initial time point. In evenly distributed time-steps, 30 µL of the reaction mix were spotted onto a 4 cm^2^ P81 filter disk (Jon Oakhill, St. Vincents Institute of Medical Research) and placed into 75 mM phosphoric acid to quench the reaction and then re-washed two times in the same solution. The filters were then rinsed with water and later with acetone, dried, and counted on the ^32^P channel of a scintillation counter as c.p.m. Control experiments were performed similarly, replacing the substrates with reaction buffer in the reaction mix. These procedures were performed in duplicate. Initial velocities were derived from a slope of a linear regression as μM of ^32^P-incorporated into substrate per minute. Reaction rates (v = *V*_0_*/E*_0_) were then calculated by dividing the slope by PKA final concentration for each experiment.

### Numerical simulations

Numerical simulations were performed using Pro-Fit 7 software (QuantumSoft) based on our kinetic model (**Supplementary Material**), to obtain the ratio of reaction rate (ξ=v_het_/v_Hom_) curves. The total enzyme concentration (E_0_) to 0.2 nM, and the total substrate concentration (S_0_) to 2 μM, resembling our experimental setup. Volume fraction of dense phase (Φ_D_) was set as an independent variable, in the range of 10^-1^ to 10^-6^. The specificity constant of the enzyme in the homogeneous system (K_SP,Hom_) was set to 1.55×10^6^ M^-1^s^-1^, which was experimentally measured for the same catalytic model in (^24^). The specificity constant of the enzyme in the dilute phase of the heterogenous system (K_SP,B_) was set to the same value, following our assumption that the reaction in the dilute phase follows the same kinetics as in the homogenous system. We then tested the effect of three variables: Partition coefficient of enzyme (K_P,E_) and substrate (K_P,S_), and the specificity constant of the enzyme in the dense phase of the heterogenous system (K_SP,D_). We varied the dimensionless K_P,E_ and K_P,S_ in five orders of magnitude, from 10^0^ to 10^5^; whereas K_SP,D_ was varied from 10^3^ to 10^7^ M^-1^s^-1^, one order of magnitude above and three below the K_SP,Hom_.

### Fluorophore labelling

For microscopy experiments, 40 μM MBD2-PKAc was labelled with 80 μM of ATTO488-NHS (Sigma-Aldrich) or CF660R dye (Biotium) in buffer 10 mM HEPES pH 8.5, 500 μL final volume reaction, at RT for 2 hours protected from light. The excess dye was removed by using CentriPure 5-Z25M (empBiotech) gel filtration columns equilibrated with buffer 10 mM Tris:Base, 150 mM NaCl, 100 μM EGTA, 10 mM MgCl_2_ pH 7.5. The sample was concentrated with AMICON 3K MWCO to 120 μL final volume, divided into 20 μL aliquots and stored at -20 °C.

For [Q_5,8_]-10 S_WT_, the labelling was performed with 40 μM of protein and 80 μM CF660R dye (Biotium) in buffer 10 mM HEPES pH 8.5, 500 μL final volume reaction, at 45 °C ON and protected from light. The excess dye was removed by cooling down the sample at 4 °C and centrifugation at 17.000 ×g for 15 minutes, after which a pellet of condensed proteins was observed. The pellet was then washed with MQ water and recovered after centrifugation. For the rest of the substrates ([Q_5,8_]-14, 20, 24, 30 and 40 S_WT_), labelling was performed in buffer 10 mM HEPES, 3 M urea, pH 8.5 at 40 °C overnight and the excess dye was removed by dialysis against MQ water using a microcentrifuge tube with a hole in the lid where a Spectra/Por2 dialysis membrane of 12-14 kDa MWCO (Spectrumlabs) was placed. After 2 days of dialysis, the formation of a pellet of condensed proteins was observed in the walls of the membranes. The content of these tubes was then heated at 90 °C for 10 minutes and the pellets were redissolved. The pellet was recovered after centrifugation at 17.000 ×g and 4°C for 15 minutes, after which the supernatant was discarded, and the pellet was washed again with MQ water and recovered after a final round of centrifugation. The pellets of all labeled substrates were resuspended in a few μL of MQ water divided into aliquots and stored at -20 °C.

Small peptides (kemptide, [Q_5,8_]-2, and [Q_5,8_]-4 S_WT_) were fluorescently labeled with CF660R dye. Labeling reactions were carried out by incubating peptides (250 μM) with CF660R (500 μM) in 100 mM sodium bicarbonate buffer (pH 8.3) at 23 °C for 2 h with shaking in a thermomixer. The reactions were quenched using reverse-phase chromatography (RPC) equilibration buffer containing trifluoroacetic acid (TFA), and excess free dye was removed by purification on a C18 reverse-phase column.

### Measurement of volume fraction of the dense phase by microscopy

To determine the total volume of the droplets in our systems with variable scaffold concentrations at 5, 10, 25, 50, and 100 μM of [Q_5,8_]-20 + [Q_5,8_]-20-p66α (9:1), 50 μL of each sample was prepared, mixed with 1 μL of Rhodamine B fluorophore to visualize the dense phase, and immediately drop-cast into a MatTek glass-bottom dish (35 mm with a 10 mm #1.5 coverslip microwell) coated with 5% (W/V) polyvinyl alcohol (PVA) to minimize wetting of the droplets. Samples were allowed to settle for 15 minutes before imaging on confocal Zeiss LSM 980 with Airyscan 2 fluorescence microscope. After settling, all droplets were confined within the first 30 μm of the sample volume, with larger droplets located near the bottom and only a few smaller droplets toward the top; the remaining volume was considered droplet-free.

Once the optical area was selected for the sample contained in the well, the lower and upper stacks were delimited. This height was fixed as the total z-axis value, and it was used for all the subsequent samples, whereas total x-y axes values were automatically defined by the acquisition set up, delimiting total sample volume. Next, a bottom-to-top 30 μm z-stack imaging in 200 steps was performed. Image stacks were analyzed using Zeiss ZEN software, and 3D segmentation was applied to reconstruct the image stacks into a 3D volume. A fluorescence intensity threshold and minimum size cutoff were applied to exclude artifacts. The summed volume of all segmented droplets was divided by the total imaged volume to calculate the droplet volume fraction (Φ_D_). Each condition was measured in triplicate, and the values were averaged.

### Fluorescence microscopy and fluorescence recovery after photobleaching (FRAP)

To perform FRAP experiments, 50 µL sample of 50 μM of [Q_5,8_]-20 + [Q_5,8_]-20-p66α (9:1) and 1 μM of CF660R-labelled [Q_5,8_]-X Substrate was mounted in a MatTek glass-bottom dish. After letting this settle for 10 minutes, an area of 20.1 μm x 20.1 μm with a droplet was selected and a small droplet region of 2.8 μm diameter (ROI) was photobleached with the confocal laser. Images were acquired using a 63x oil objective (Plan-Apochromat NA = 1.4) and a 639 nm laser beam at 0.2% laser power at room temperature and detected between 650 nm and 690 nm using a PMT detector measuring with a gain of 750 V and an offset of 0%. Pre- and post-bleach images were acquired at 0.5-second intervals for a time span of 100 seconds. Fluorescence recovery was monitored at a resolution of 512 × 512 pixels, 16-bit depth, with a final pixel size of 0.043 µm/pixel and measured over time. The recovery curves were generated from the average fluorescence intensity within the ROI of three independent acquisitions per Substrate. Recovery Time (τ) for each curve was determined by fitting the average data to a simple exponential recovery model:

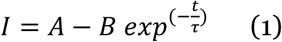

The mobile fraction was calculated as:

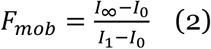

The apparent diffusion coefficient from FRAP was calculated according to:^27^

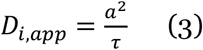

where “a” is the radius of the photobleached regions (ROI).

### Confocal fluorescence microscopy to measure partition coefficients

Partitioning of substrates of different lengths into [Q_5,8_]-20 condensates was determined using the single-photon counting confocal microscope Luminosa (PicoQuant). Substrates were labeled with the CF660R dye to enable visualization of their partitioning into the scaffold. For sample preparation, 1 µM of CF660R-labeled substrate or enzyme was mixed with the scaffold [Q_5,8_]-20 + [Q_5,8_]-20-p66α (9:1) at a final scaffold concentration of 50 µM and drop-cast onto PVA-coated MatTek glass dishes. Samples were left to equilibrate for several minutes before imaging at room temperature.

The Luminosa system is based on an inverted Olympus IX73 microscope equipped with a 60× PlanApochromat water-immersion objective (NA = 1.2, Olympus) and single-photon avalanche diode (SPAD) detectors (PicoQuant, Pi Imaging). Fluorescence was excited using a 640 nm pulsed diode laser (LDH-D-C-640S) with a repetition rate of 40 MHz and laser powers ranging from 54 to 296 nW, depending on the sample fluorescence intensity. The laser power was selected to ensure stable fluorescence emission within condensates, while remaining sufficiently high to reliably detect fluorescence counts in the dilute phase. Fluorescence emission was collected through a 485 nm dichroic mirror (Chroma), a 655–725 nm bandpass filter (AHF Analysentechnik), and a 102 µm pinhole.

Before acquiring point measurements, condensates were imaged at their central focal plane, close to the sample surface, in order to identify condensate positions (dense phase). Both a transmission image using white light and a camera, as well as a fluorescence image using the laser and point detector, were acquired. An additional image was recorded approximately 30 µm above this plane to confirm the absence of condensates (dilute phase). Imaging was performed over an area of 203 × 203 µm^2^ with a pixel dwell time of 30 µs and a pixel resolution of 198 nm. The Q20S scaffold in the absence of fluorescent substrate was measured at comparable laser powers, yielding background photon counts below two photons, indicating negligible autofluorescence. Subsequently, ten individual point measurements were acquired for both the dense and dilute phases, with an acquisition time of 20 s per point.

Point measurement data were analyzed by calculating the average photon count rate (kilocounts per second, kcps) for each measurement, followed by the determination of the mean kcps and corresponding standard deviation across all ten measurements. The partition coefficient was determined as the ratio of the mean kcps in the dense phase to that in the dilute phase, with the propagated standard error.

## Results

To study how biomolecular condensates affect kinase phosphorylation kinetics, we established a modular model system based on the resilin-like peptide condensates and protein kinase A (PKA). The kinase and its cognate substrate were recruited to the condensate as clients by distinct mechanisms: the kinase via a protein-protein interaction motif linked to an RLP and the substrate through direct fusion with RLP-repeat regions (**Fig. 1A**). To avoid off target phosphorylation, we changed all serine residues from the consensus RLP into glutamine residues resulting in the repeat sequence GRGDQPYQ.

**Figure 1.**
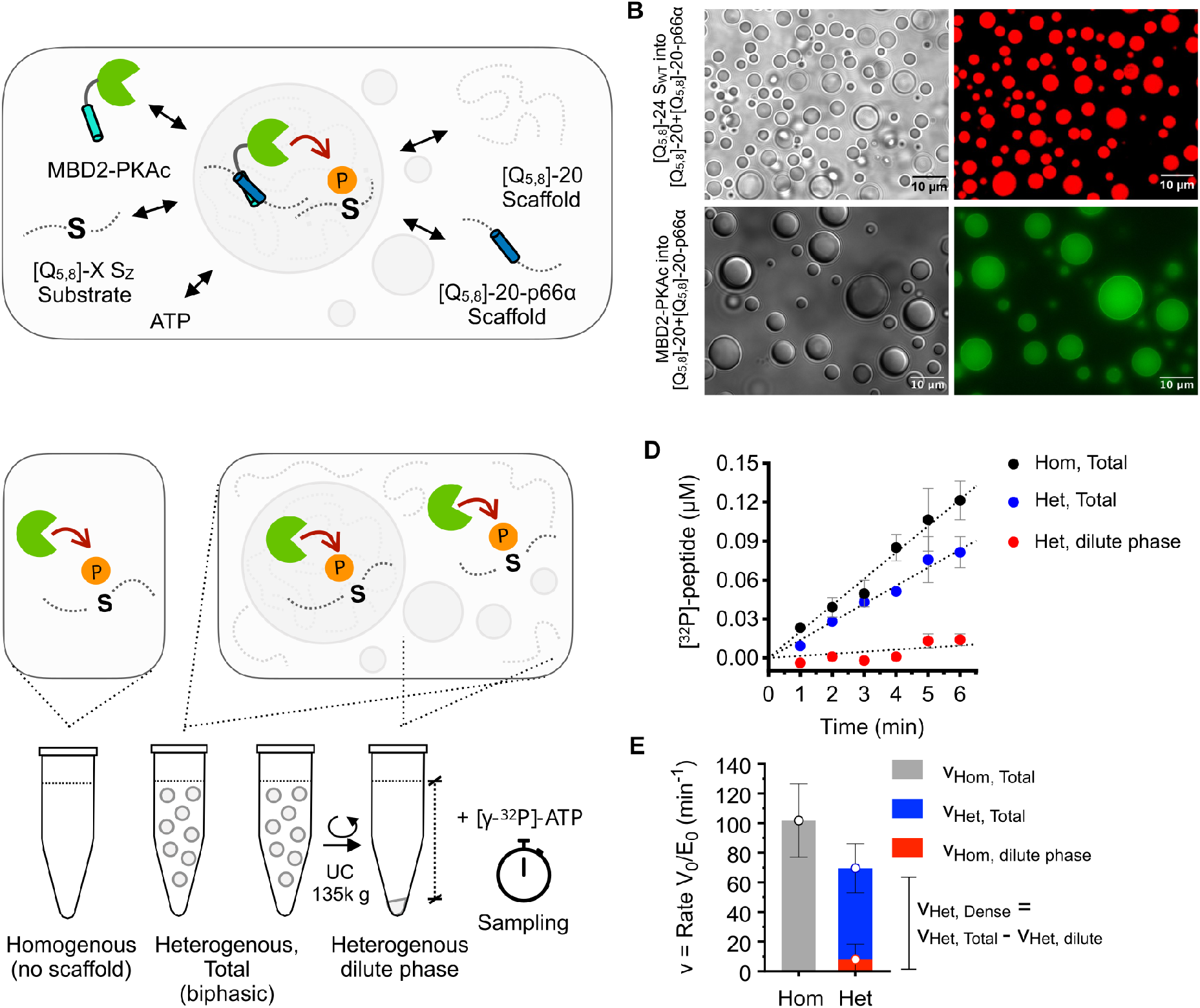
A modular model system for PKA phosphorylation in condensates. **A)** Schematic of condensates model system: The condensate scaffold consisted of [Q_5,8_]-20 with 10% [Q_5,8_]-20-p66α to increase recruitment of the kinase through coiled-coil interactions with MBD2-PKAc. Kinase and substrate are recruited as sub-stoichiometric clients. B) DIC and confocal fluorescence microscopy showed spherical condensates that concentrate fluorophore labelled substrate (upper) and enzyme (lower). **C)** Procedure for kinetic experiments, where dense and dilute phases are separated by ultra-centrifugation and analyzed as kinetic experiments.^20^ **D)** Time course of ^32^P-phosphorylated [Q_5,8_]-24S_WT_ substrate (2 μM) in homogeneous solution and a biphasic (50 μM scaffold) and dilute phase samples. Each measurement was done in duplicate (n=2) and error bars correspond to the standard deviation. **E)** Decomposition of phosphorylation reaction rates (v_Hom_, v_Het,Total_: v_dil_ and v_D_) of [Q_5,8_]-24S_WT_ reveals that the condensates are overall inhibitory. The height of the bar represents the calculated in vitro phosphorylation reaction rate (n=1) for each condition, and the error bars correspond to the propagated standard error of the rates.

The host condensate [Q_5,8_]-20 forms dynamic spherical condensates with a C_sat_ of 0.54 ± 0.014 mg/mL in agreement with previous studies.^28^ The consensus substrate peptide of PKA (kemptide, S_WT_: LRRASL) was introduced into resilin repeat proteins with 24 repeats ([Q_5,8_]-24 S_WT_) to allow it to be distinguished from the scaffold protein by gel electrophoresis. Fluorescently labeled substrates are strongly recruited into the condensate (**Fig. 1B**), and at the concentrations used this substrate recruitment did not alter the phase behavior properties of the host condensate (**Supplementary Fig. 2**). The catalytic domain of PKA alone did not show strong partitioning into the condensates.

Therefore, we created a modified scaffold where the coiled-coil domain p66α is introduced at the center. P66α forms a nanomolar affinity complex with MBD2,^29^ which allows recruitment of PKA fused to the coiled-coil of MBD2. When the host condensate contains 10% of the p66α containing RLP, the kinase is strongly recruited throughout the condensate (**Fig. 1B**). In summary, this creates a modular system where the kinase and substrate primarily encounter each other inside the host condensate.

We used ^32^P radio assays to assess the phosphorylation kinetics comparing reactions with and without condensates (**Fig. 1C**). In parallel with the biphasic sample, an identical biphasic sample is cleared by ultra-centrifugation to remove all condensates and isolate the dilute phase. We thus recorded parallel kinetic assays for a homogeneous sample without condensates, a biphasic sample and a dilute sample corresponding to the dilute phase (**Fig. 1D**). This allowed us to isolate the contributions from the dense phase by subtracting the contribution of the dilute phase from that of the biphasic sample (**Fig. 1E**).^20^ For [Q_5,8_]-24 S_WT_ under biphasic conditions, this analysis showed that the dilute phase only contributes minimally to the reaction due to the strong partitioning of substrate and enzymes into the condensate. It is also clear that the presence of the condensates retards the progress of the phosphorylation reaction, as also confirmed by Phos-Tag gel^30^ (**Supplementary Fig. 3**). This is contrary to the effect predicted by mass action, where co-partitioning of enzyme and substrate is expected to lead to increased catalytic rates. This suggested that there is an inhibitory effect of the condensate environment.

### Probing the structure and dynamics of PKA in [Q_5,8_]-20 condensates

The condensate environment is more hydrophobic than the dilute phase, which can affect the structure of client proteins, e.g. by transient local unfolding^31^ or favoring open conformations.^7^ Hydrogen-deuterium mass spectrometry has emerged as one of the few techniques that can assess the folding and dynamics of proteins inside condensates^31,32^. We thus performed hydrogen-deuterium exchange on MBD2-PKAc in the presence and absence of [Q_5,8_]-20 condensates (**Fig. 2A**). Whereas MBD2-PKAc gave high peptide coverage in the absence of condensates (**Fig. 2B**), the presence of condensates suppressed peptide coverage in PKA. To maximize the signal from MBD2-PKAc, we thus increased the concentration of the client to 20 μM, i.e. a 1:10 ratio between client and scaffold. Previous HDX studies of phase separating proteins have either isolated the dense phase by centrifugation^31^ or analyzed a biphasic mixture.^32^ We found that pelleting of the condensates had a strong negative effect on peptide recovery, and thus proceeded with analysis of a biphasic system. The combination of a high partitioning coefficient and a volume fraction (6 %) means that (76 %) of the enzyme is in the dense phase. Under these conditions, we detected 105 peptides, resulting in 76.8% sequence coverage of MBD2-PKAc **(Supplementary Fig. 4**). These conditions correspond to a dynamic partitioning equilibrium dominated by the dense phase.

**Figure 2.**
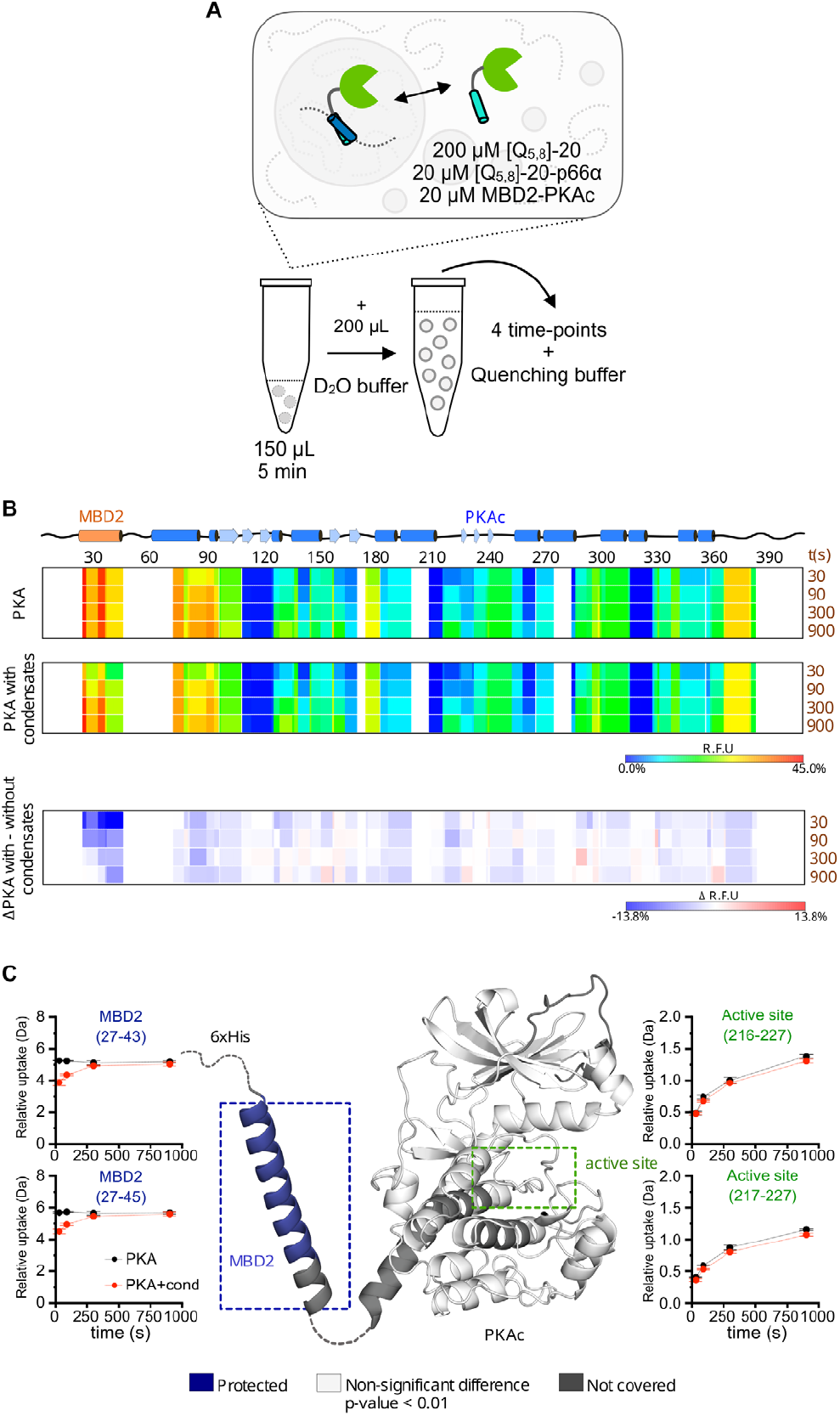
Hydrogen-deuterium exchange reveals that PKA is structurally intact in condensates. **A)** Experimental setup for hydrogen exchange experiment: MBD2-PKAc was recruited as a sub-stoichiometric client to [Q_5,8_]-20 condensates followed by dilution in D_2_O buffer and subsequent quenching at pH 2.2. **B)** Hydrogen exchange mapped onto peptide chain for the PKA alone (top), in the presence of condensate (middle) and the difference (bottom). Segments in white indicate lack of peptide coverage. **C)** Mapping of coverage and significant changes in hydrogen exchange on a structural model of the PKA catalytic domain (PKA: 1APM). Deuterium uptake profile of 2 peptides corresponding to the coiled-coil domain of MBD2 (blue) and 2 peptides at the active site region (green). Red line and dots represent uptake of MBD2-PKA without condensates and black line and dots represent uptake of MBD2-PKA with condensates. Deuterium measurement was repeated in parallel experiments (w. condensates n=4, reference n=5) and the error bars correspond to standard error of the mean.

Comparison of the deuterium uptake in MBD2-PKAc indicates that the structure and dynamics of the kinase domain are broadly preserved inside the condensate (**Fig. 2B**). Only one region in the client protein shows significant changes in deuterium at early time points (**Fig. 2C**). This region corresponds to the coiled-coil domain of MBD2, which is used to recruit the kinase to condensate by binding to the matching coiled-coil domain in [Q_5,8_]-20 p66α. The observed protection is consistent with effective interaction upon binding. In contrast, peptides around the catalytic site show virtually indistinguishable deuterium uptake, suggesting that the dynamics of the domain are unchanged (**Fig. 2C**). In conclusion, the HDX experiments indicate that the reduced phosphorylation rates are not caused by a structural change of the kinase inside [Q_5,8_]-20 condensates.

### Numerical simulations of phosphorylation in biphasic systems

Next, we explored a biphasic system, where the condensates affect both the rate constants and establish a partitioning equilibrium of enzymes and substrates. We derived an analytical equation describing an enzymatic reaction in a biphasic system (**Supplementary Text**). In our kinetic model, we consider a system in which both enzyme and substrate partition between the two phases, with characteristic partitioning coefficients, K_P,E_ and K_P,S_ (**Fig. 3A**). The catalytic efficiency is expressed as specificity constants, K_SP,dil_ and K_SP,D_, which correspond to k_cat_/K_M_ in the classical Michaelis-Menten equation, but provides more robust fitting of enzyme kinetic data when saturation cannot be conveniently reached.^33^ Throughout the following, the rate for a biphasic system is compared to a homogeneous system with similar concentrations of enzyme and substrate, and where the rate constant of the dilute phase equals that of the homogeneous solution. The resulting ratio (ξ = v_Het_/v_Hom_) thus quantifies the effect of phase separation, where ξ >1 indicates acceleration and ξ < 1 indicates inhibition.

To explore the interplay between mass action and an inhibitory dense phase environment, we simulated ξ for five orders of magnitude of variation of K_P,S_, K_P,E_, K_SP,D_ and the volume fraction of condensates Φ_D_ (**Fig. 3B**). The rate constants of the dilute phase are fixed at experimental values for the phosphorylation of kemptide by PKA at K_SP_ = 1.55 × 10^6^ M^-1^ s^-1^.^24^ The effect of mass action is seen clearly in the simulation using similar rate constants in the dense and dilute phase (K_SP,D_ = 10^6^ M^-1^s^-1^). The rate enhancement forms a bell shape curve as a function of the Φ_D._ The initial increase occurs as the fraction of enzyme and substrate concentrated in the dense phase increases, whereas the decline occurs when the dense phase concentration declines due to dilution. The rate enhancement in such systems can exceed 100-fold and can only retard the reaction if enzyme and substrate partition into different phases. The optimal enhancement is reached at lower Φ_D_ for strongly partitioning systems in line with previous simulations.^34^ When the catalytic rate in the dense phase is increased to 10^7^ M^-1^s^-1^, the curves retain a broadly similar shape, but with an elevated enhancement.

**Figure 3.**
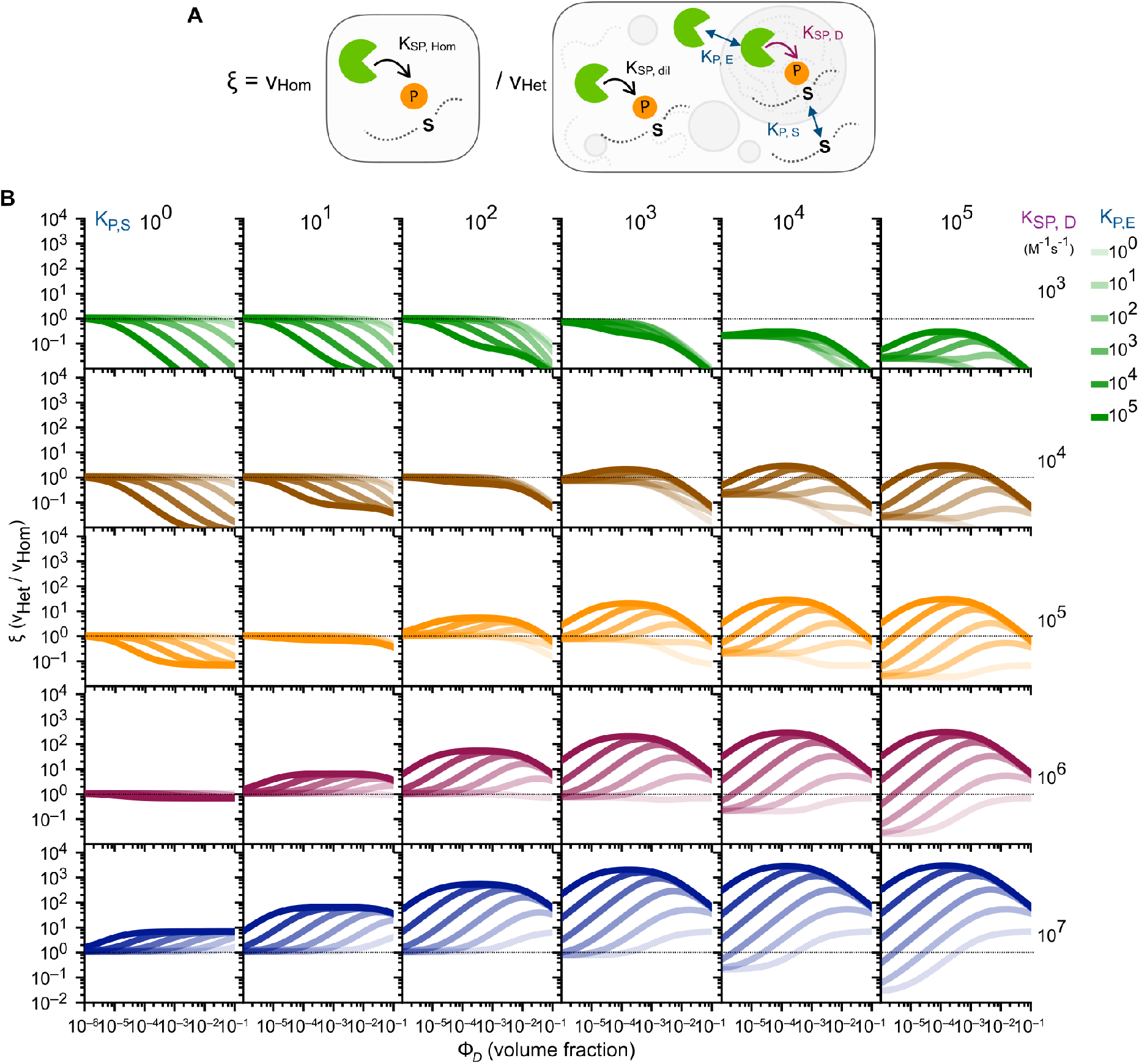
Numerical simulations of enzyme catalysis in a phase separated system. **A)** The numerical simulations compare the phosphorylation rate in homogeneous solution (v_Hom_) with a phase separated system (v_Het_) where enzyme and substrate distributing in the two phases with variable partitioning constants (K_P,S_ and K_P,E_). The specificity constant is identical in the dilute and homogeneous phase (K_SP,dil_ and K_SP,Hom_), but varied in the dense phase K_SP,D_. **B)** Rate of the heterogeneous system relative to that of an identical homogeneous system as a function of volume fraction of condensates as a function of K_P,S_ (horizontal), K_P,E_ (color density) and K_SP,D_ (vertical). Simulations are based on the equations developed in **Supplementary Materials**.

Simulations with a lower K_SP,D_ (10^6^-10^6^ M^-1^s^-1^) show a varied dependency on Φ_D_ ranging from a stimulatory bell shape to a monotonous inhibition by condensates (**Fig. 3B**). In between these extremes there are intermediate scenarios where the condensate accelerates reactions at low Φ_D_ but becomes inhibitory at low high Φ_D_. Such behavior is characteristic of a trade-off between acceleration by mass action and an inhibitory intra-condensate environment. In total, these simulations show that a broad range of behaviors can be expected for two-phase enzyme substrate systems, and that the same condensate can be both inhibitory and stimulatory depending on the parameters of the biphasic system. From an experimental point of view, this suggests that it is instructive to study enzyme reactions across a range of Φ_D_ values.

### Phosphorylation kinetics of different substrate lengths and volume fractions

Next, we sought to modulate the partitioning of the substrate into the condensates without affecting the recognition of the substrate motif by the kinase. The repeat protein design provides a unique opportunity to gradually tune partitioning, without relying on stereospecific interactions, by changing the number of repeats. We thus made kemptide substrates containing 2, 4, 10, 14, 20, 24, 30, and 40 repeats ([Q_5,8_]-x S_WT_), in addition to kemptide alone, creating substrates spanning from 1 to 40 kDa (**Fig. 4A**). Initially, we compared the phosphorylation kinetics of these substrates in a homogeneous solution (**Fig. 4B, Supplementary Table 3**). Substrates with between 10 and 30 repeats have similar phosphorylation rates, consistent with those previously determined for kemptide in other fusion proteins.^24,35^ [Q_5,8_]-40 S_WT_ has noticeably slower kinetics that are approximately 3-fold slower than expected. The concentration used in the enzymatic assay (2 µM) is above the C_sat_ expected for such a long RLP,^28^ and accordingly we observe condensates even in the absence of scaffold. We thus excluded this substrate from further comparisons. For substrates with 4 repeats or fewer, we observed significantly lower phosphorylation rates using the ^32^P assay. This is likely due to a known systematic artifact in ^32^P kinase assays, where short peptides bind poorly to phospho-cellulose filters.^36^ In the following, we will compare the ratio of phosphorylation rates, which should largely cancel this artifact.

**Figure 4.**
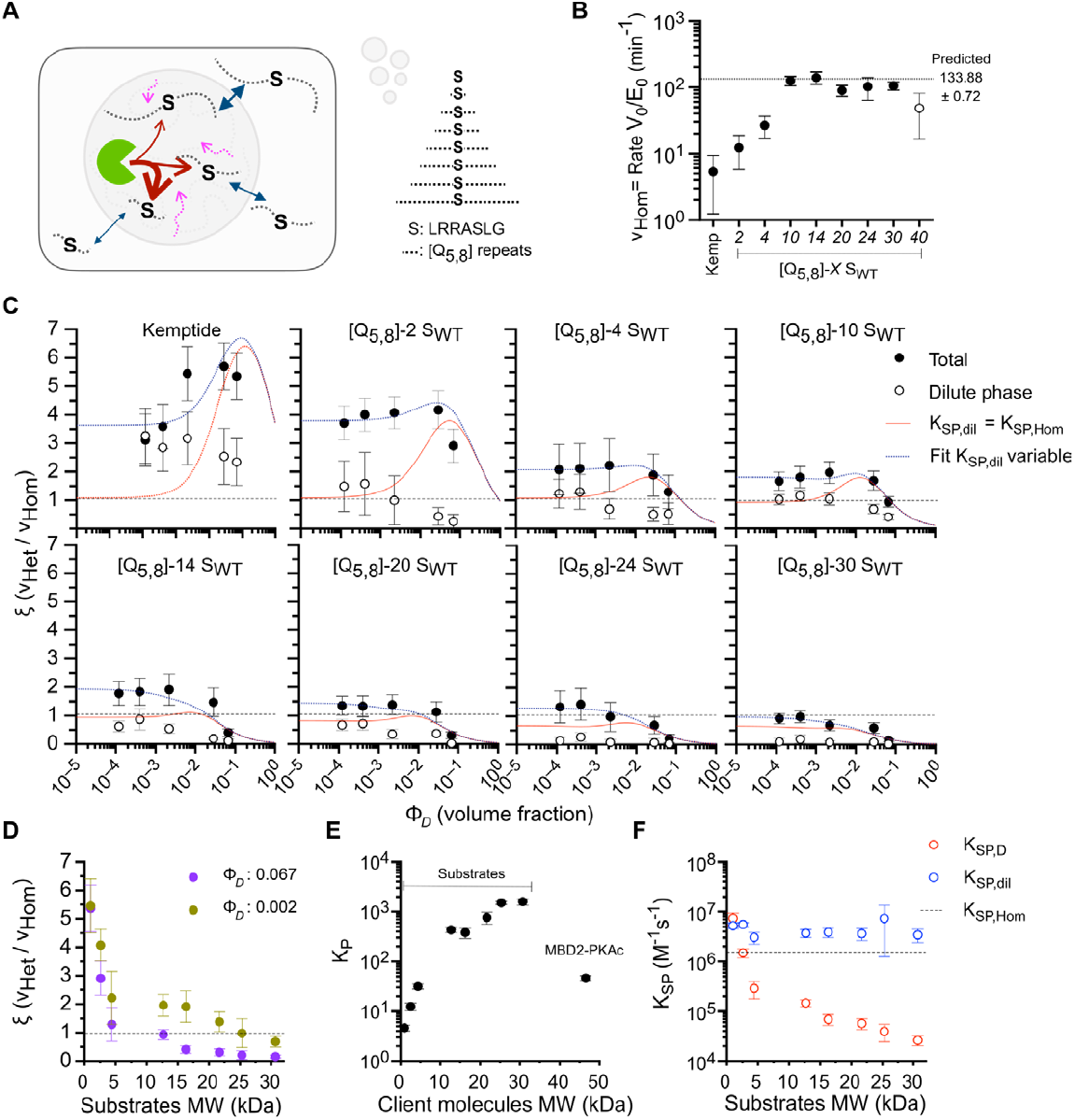
Substrate length dependence of phosphorylation. **A)** Variation of substrate length by appending 0-40 eight residue repeats to the kemptide substrate motif to vary diffusion and partitioning. The substrate motif is always placed in the middle of the sequence. **B)** Observed in vitro phosphorylation reaction rates in homogeneous solution from ^32^P kinase assay for different substrates. The lower rates for short constructs are likely a consequence of poor binding of short peptides to filter paper. The longest substrate, [Q_5,8_]-40S_WT_, forms condensates at the concentration used and is excluded from further experiments. _32_P kinase assays were done in duplicate (n=2) and one phosphorylation rate value was calculated for each substrate. The error bars correspond to the propagated standard error of the rates. **C)** Ratio of reaction rates (ξ) in the presence of condensates relative to homogeneous conditions (Total: v_Het,Tot_/v_Hom_, black circles, and Dilute phase: v_Het,Tot_/v_Hom_, empty circles) as function of volume fraction of the dense phase. The kinetics for each condition was determined as outlined in Fig. 1 C-E. Error bars correspond to the propagated standard error of the rates. Lines represent a fit to the model defined in the **Supplementary Text.** with partitioning coefficients constrained to values in Panel E. In the first fit (red), K_SP,dil_ was constrained to K_SP,Hom,_ in the second fit K_SP,D_ was constrained to the value from the first fit and K_SP,dil_ was allowed to vary. **D)** Values from Panel C replotted as a function of molecular weight of the substrate for scaffold concentrations of 25 μM (Φ_D_ = 0.002) and 100 μM (Φ_D_ = 0.067). **E)** Partitioning coefficient of substrates and enzyme measured by fluorescence confocal microscopy (n=10). Error bars represent the propagated standard errors. **F)** Specificity constants from the dense and dilute phase from fits in Panel C. Error bar represents the standard error of the fitting. The line represents K_SP,Hom_ for kemptide (1.55 μM^-1^s^-1^). ^24^

Inspired by the simulations above, we performed ^32^P kinase assays in the presence of condensates at different volume fractions (**Supplementary Fig. 5, Supplementary Table 2**). We varied the concentration of [Q_5,8_]-20 scaffold from 5 µM to 100 µM, which resulted in condensate volume fractions ranging from 0.0001 to 0.0669 (**Supplementary Table 2**) as determined by confocal microscopy. The dynamic range of the series is limited at the lower end by a desire to maintain a comparable condensate across the series, which suggests that the substrate should be a minor component. At the higher end, it is limited by the maximum density of condensates that remain suspended during the enzymatic assay.

We compared the ratio of phosphorylation rates (ξ) across different volume fractions of condensate for all the substrates (**Fig. 4C**). Each curve is consistent with an initial increase at low volume fractions of condensate followed by a decay at high volume fractions. For long substrates this takes the form of a plateau and for short substrates an apex at intermediate volume fractions, where the optimal volume fraction decreases with substrate length. The contribution of the dilute phase can be isolated by removing the dense phase by centrifugation and performing a parallel assay with only the dilute phase. For the longest substrates, the contribution of the dilute phase is negligible, and the total reaction is dominated by the reaction occurring in the dense phase. The contribution of the dilute phase increases for short substrates, likely due to a weaker partitioning of the substrate into the condensate. Surprisingly for kemptide, the dilute phase has a higher catalytic rate than the sample without condensates (**Supplementary Fig. 6**). When compared at similar volume fractions, the rate enhancement decreases uniformly with the molecular mass of the substrate (**Fig. 4D**). At low Φ_D_ values, it decreases to a value near the homogeneous rate, whereas at high Φ_D_ values it is depressed to much lower values.

Directly fitting the above data to extract kinetic rate constants is an ill-posed problem due to the many variables in the equations describing the two-phase system. To constrain the fitting, we measured partitioning of all substrates and kinases by fluorescence microscopy. As expected, the substrates partition much more strongly into condensates with increasing length (**Fig. 4E, Supplementary Table 3**), with kemptide having a K_P_ of ∼5 and [Q_5,8_]-30 S_WT_ of ∼1590. We fitted the data in Fig. 4C with the kinetic model described in **Supplementary Text**, where the values for K_P_ and Φ_D_ were constrained to the values determined by microscopy. Initially, the K_SP,dil_ was constrained to the rate in the absence of condensates (K_SP,Hom_). This model describes longest substrates well, including the decrease in phosphorylation at high volume fractions. For the shorter substrates, the model fails to account for the increased rates at low volume fractions, as constraining K_SP,dil_ to K_SP,Hom_ forces the curve to asymptotically approach ξ = 1. Moreover, the width of the peak of elevated v_Het_ values suggests that mass action alone is insufficient to account for this. Next, we allowed the values K_SP,dil_ to vary, while constraining K_SP,D_ to the value from the previous fits. This allowed the fits to converge for all substrates (**Fig. 4C, Supplementary Table 4**). Independent fits to data sets for different substrate lengths converge on a K_SP,dil_ that is approximately three times the rate in the absence of condensates (**Fig. 4F**). This suggests the presence of a species at equilibrium with the dense phase that is responsible for the rate acceleration, which is consistent with observations in other engineered enzyme-condensate systems.^20,37^ The enzymatic rate constant in the dense phase (K_SP,D_) shows a monotonous decrease with substrate length (**Fig. 4F**). For the longest substrates, the rate constant is decreased a hundred-fold relative to the rate constant in the absence of condensate. This decrease cancels the effect of concentrating enzyme and substrate into the same compartment.

### Diffusion of substrates in condensates

Next, we measured the diffusion of PKA substrates as clients in [Q_5,8_]-20 condensates. The percolated networks of biomolecular condensates give rise to probe-size-dependent viscosity that deviates from Stokes-Einstein behavior. Small molecules diffuse at speeds similar to those in dilute solutions, whereas large macromolecules diffuse according to the viscosity measured by macroscopic techniques.^38,39^ Our substrates fall in the transition between these regimes. We performed FRAP experiments using CF660-labelled substrates recruited as clients into a [Q_5,8_]-20 condensate (**Fig. 5A, Supplementary Fig. 7**). All substrates showed near full recovery within a few minutes, confirming that the condensates remain dynamic in the experimental timeframe used in kinase assays. The FRAP recovery was fitted to extract the mobile fraction (**Fig. 5B**) and the recovery time constant (**Fig. 5C**) for each substrate (**Supplementary Table 5**). The shortest substrates experienced full recovery whereas the mobile fraction decreased to about 80% for the longer substrates, suggesting the presence of a small fraction of immobile species. The recovery time constant showed a dramatic change for the shorter substrates but reached an apparent plateau for substrates longer than [Q_5,8_]-14.

**Figure 5.**
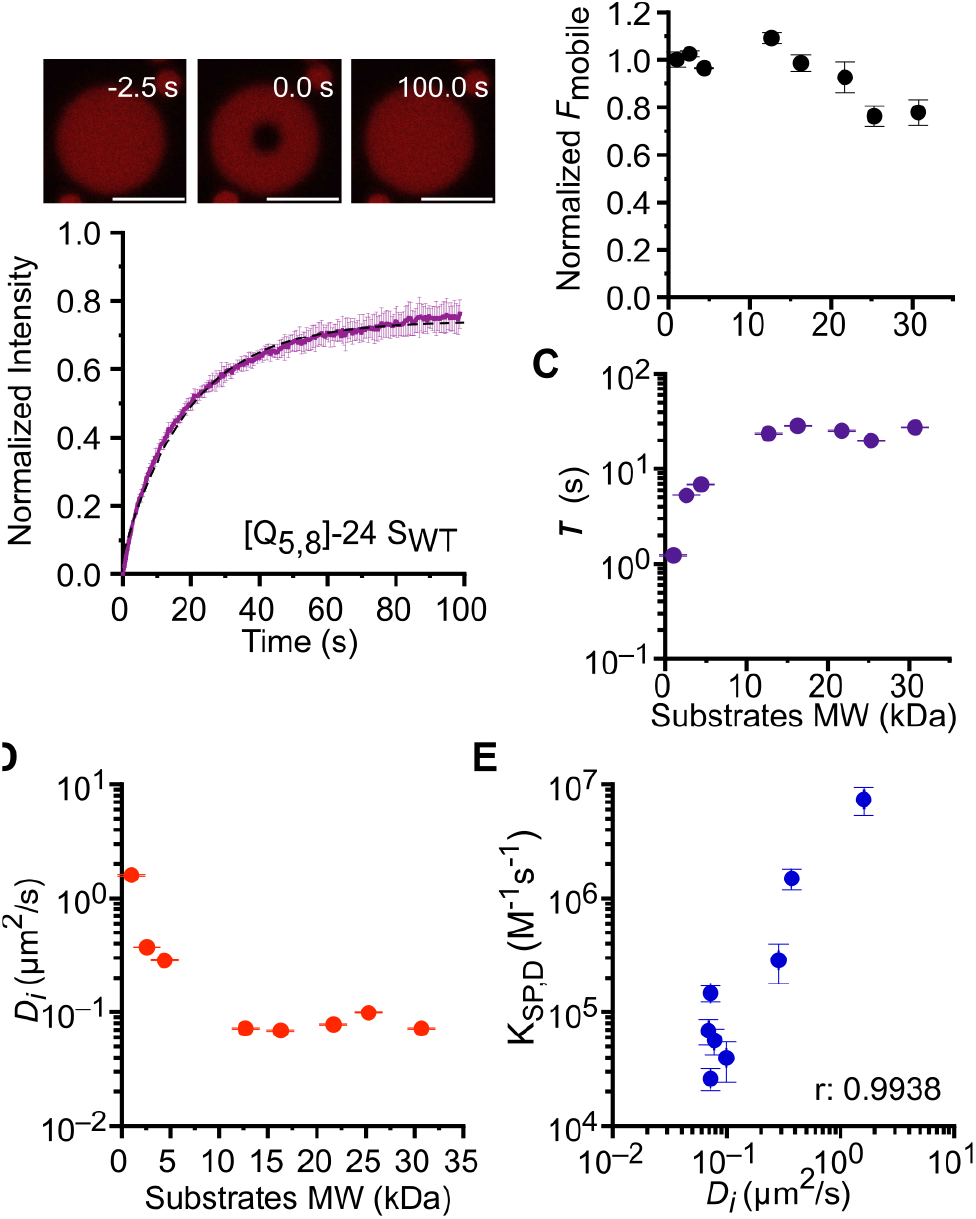
Diffusion dependence of phosphorylation rates. **A)** FRAP of 1 μM CF660R-[Q_5,8_]-24 S_WT_ recruited as client into 50 μM [Q_5,8_]-20 (scale bars correspond to 10 μm). Three independent FRAP-determinations (n=3) were performed per substrate and averaged. Dashed line represents the fit to a simple exponential recovery model. **B, C)** Fitted parameters from the model used including the mobile fraction (B) and recovery time (C). **D)** Diffusion coefficient calculated from recovery times in C based on *D*_*i,app*_= a^2^/τ. Error bars correspond to the standard error of the mean. **E)** Correlation between diffusion coefficient of substrates and specificity constant in the dense phase.

We extracted apparent diffusion coefficients for the substrates (**Fig. 5D**). The diffusion coefficients of the long substrates are approximately 1000-fold lower than expected from similar proteins in dilute solution, but similar in magnitude to values for other IDPs in condensates.^23^ The diffusion coefficients followed a trend similar to the recovery times, gradually decreasing with substrate length up to [Q_5,8_]-14, whereafter they plateaued at similar values. The plateau is similar to what was observed for the diffusion of other IDPs in FUS condensates, where little length dependence was observed.^40^

The diffusion coefficients of the substrates are highly (r =0.99) correlated with the rate of the phosphorylation reaction (**Fig. 5E**). The decrease in K_SP,D_ for the longest substrates cannot be readily explained by the diffusion constants extracted from the time constant of FRAP recovery. However, the mobile fraction decreases slightly with substrate length for the longer substrates, indicating the presence of a long-lived pool of immobile substrates that contributes to depress the rate of phosphorylation. In total, this suggests that the slow diffusion of substrates in the condensate is a key factor in decreasing phosphorylation rates.

### Origin of diffusion-limitations

There are at least two distinct ways in which slow substrate diffusion can retard an enzymatic reaction. In a classical diffusion-limited enzyme reaction, diffusion controls the frequency of encounters between the active site of the enzyme and the substrate (**Fig. 6A**). In vitro studies using viscogens have shown that solvent viscosity can retard especially fast enzymes limited by k_cat_.^41^ Alternatively, in phase separated systems the inhomogeneous distribution of enzymes and slow diffusion can lead to mass transport limitations. Highly concentrated enzymes constitute a localized sink of substrates, where the condensate experiences a non-equilibrium depression of substrate concentrations due to the constant enzymatic turnover (**Fig. 6B**).^8^

**Figure 6.**
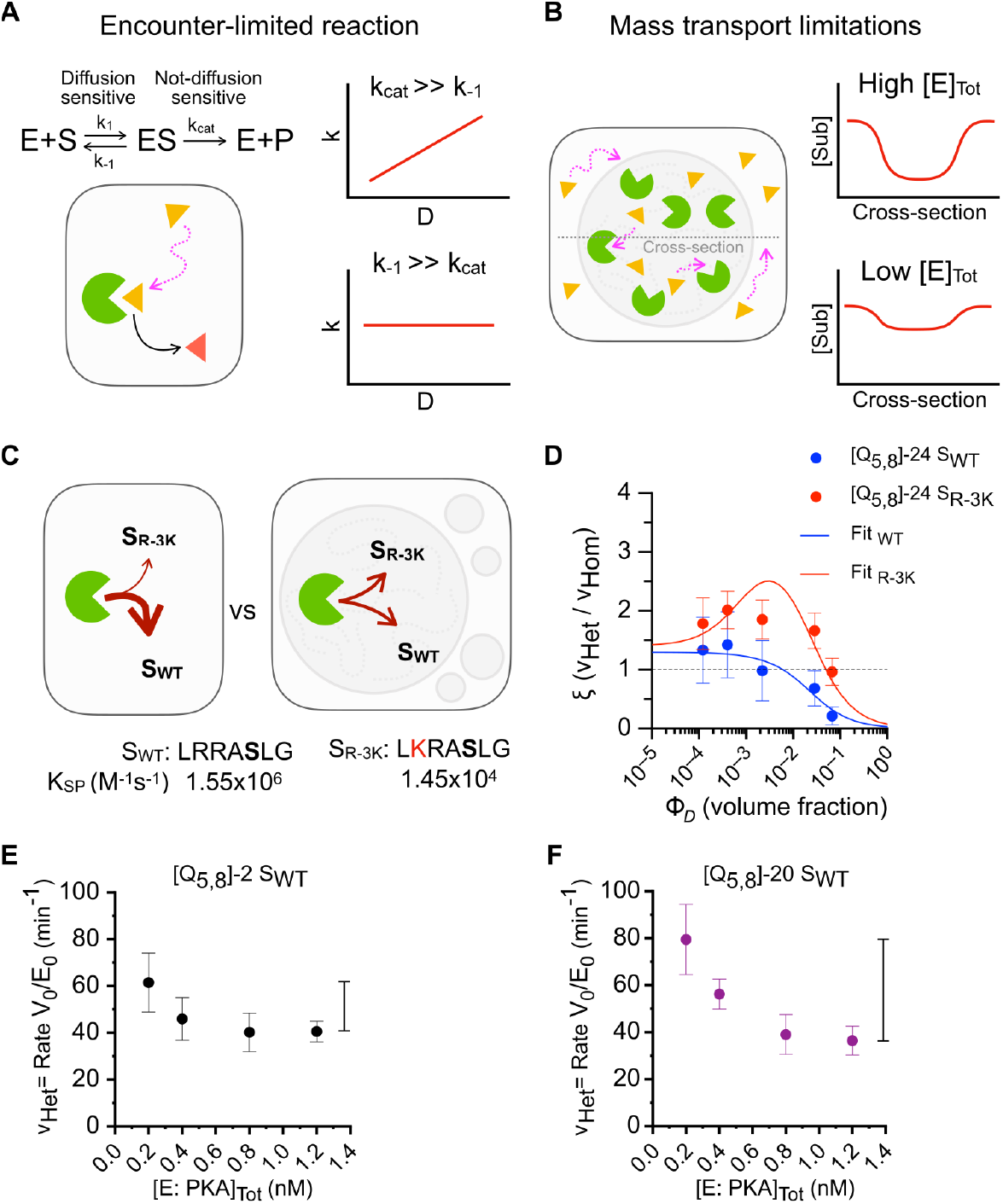
Diffusion-limitations arise from mass transport limitations. **A)** Mechanistic models for diffusion-limitation in condensates. Classical diffusion-limitations as observed as viscosity dependence in homogeneous solution occur primarily for very fast enzymes, where the limiting step in the reaction is the association of enzyme and substrate. **B)** A high concentration of enzymes in the dense phase can deplete the substrate concentration locally. Unlike reactions that are limited by the association rates, mass transport limitations should be more severe for more concentrated enzymes. **C)** Schematics showing that the phosphorylation kinetics is reduced 100-fold by a point mutation in the substrate motif (S_R-3K_) whereas inside condensates this could change. **D)** Comparison between the ratio of phosphorylation reaction rates (ξ) in the presence of condensates relative to homogeneous conditions for substrates S_WT_ and S_R-3K_ embedded in [Q_5,8_]-24 chain. Lines represent the fit of [Q_5,8_]-24 S_WT_ as outlined in Fig. 4 C (blue), whereas for [Q_5,8_]-24S_R-3K_ there were no constraints and K_SP,D_ and K_SP,dil_ were allowed to vary. **E, F)** Enzyme concentration dependency of phosphorylation reaction rates for [Q_5,8_]-2 S_WT_ (D) and [Q_5,8_]-20 S_WT_ (E) in 50 μM [Q_5,8_]-20 scaffold (Φ_D_= 0.028). The kinetics for each condition were determined as outlined in Fig. 1 C-E. Error bars correspond to the propagated standard error of the rates.

To distinguish between these mechanisms, we probed the effect of the intrinsic rate constants on the kinetic enhancement of the condensates. The kinetics of PKA phosphorylation of kemptide is similar to enzymes previously found to exhibit viscosity dependence.^41–43^ Classical viscosity dependence suggests that a PKA substrate with slower kinetics would not experience the same retardation in the dense phase. The phosphorylation kinetics of PKA is highly dependent on the recognition motif near the phospho-acceptor.^44^ A mutation (S_R-3K_: L*K*RASL) from Arg to Lys (S_R-3K_: L*K*RASL) reduces K_SP_ by ∼100-fold, mainly through an elevated K_M_ (**Fig. 6C**).^24^ Furthermore, the broadening of substrate usage observed in other model systems^21^ suggests that the condensates should enhance the S_R-3K_ more than S_WT_. We therefore investigated the phosphorylation kinetics of [Q_5,8_]-24 S_R-3K_ as a function of dense phase concentration of [Q_5,8_]-20 condensate as above (**Fig. 6D, Supplementary Fig. 8, Supplementary Table 6**). The phosphorylation rate S_R-3K_ follows broadly the same pattern as seen for S_WT,_ albeit with an elevated plateau at low volume fractions. Fitting of the S_R-3K_ suggested that the data can be accounted for by ∼2-fold higher K_SP,dil_ and a similar decrease in K_SP,D_ as for S_WT_. This indicates a modest increase relative to the S_WT_, but not to the same extent as seen previously.

In contrast to diffusion-limitation on the association step, mass-transport limitation is a meso-scale phenomenon that arises due to the inhomogeneous distribution of enzymes. If enzymes are clustered sufficiently densely, the rate at which substrate is consumed can exceed its replenishment by diffusion. In chemical engineering, mass-transport limitations are often estimated by the Damköhler number, Da = τ_D_/τ_R_, which compares the time scales of diffusion and chemical reactions and was recently extended to condensates.^37^ For the substrates in the dense phase, τ_D_ can be estimated from the characteristic radius (r = ∼2 µm) using τ^D^ = r^2^/D_i_, yielding values ranging from 2.5 to 56 s. The timescale of reaction can be estimated assuming pseudo-first order kinetics as τ_R_ = (K_SP_ [E]_D_)^-1^. Using an [E]_D_ = ∼50 nM and K_SP_ from homogeneous solutions, we estimate Damköhler numbers of ∼0.2, ∼0.8, and ∼4 for kemptide, [Q_5,8_]-2 S_WT_, and [Q_5,8_] 10-30 S_WT_, respectively. These numbers are crude approximations but nonetheless suggest an increase from moderate to severe mass-transport limitations with increasing lengths of substrates.

To evaluate mass-transport limitations experimentally, we varied the enzyme concentration of the phosphorylation assay. As substrate consumption depends on the enzyme concentration while diffusion does not, mass-transport limitations will increase with increased enzyme concentration, in contrast to diffusion-limits acting on the association (**Fig. 6B**). We thus recorded phosphorylation kinetics in condensates as a function of PKA concentration for substrate with fast diffusion [Q_5,8_]-2 S_WT_ (**Fig. 6E**) and slow diffusion [Q_5,8_]-20 S_WT_ (**Fig. 6F, Supplementary Fig. 9, Supplementary Table 7**). Due to the differing detection efficiencies of phospho-peptides (**Fig. 4B**), the absolute values from these experiments are not directly comparable; however, the concentration dependence should be. For the short substrate, there is a modest reduction with increasing enzyme concentration, but the effect is still within error. For the long substrate, the enzyme-normalized rate is 2-fold higher at lower concentrations. In combination, these experiments suggest that the dependence on diffusion coefficients is mostly due to mass transport limitations rather than effects on the formation of enzyme-substrate complexes.

## Discussion

We have shown that the presence of condensate forming proteins can either enhance or depress the rate of a phosphorylation reaction, depending on the volume fraction. The enhancement seems to arise from the dilute phase, whereas the condensate itself is inhibitory to the reaction in spite of a strong colocalization of enzyme and substrate. The inhibition is not due to changes in the structure and dynamics of the enzyme, but rather arises from the slow diffusion of substrates, particularly those embedded in long chains, in a manner that seems to invoke mass transport limitations rather than a reduced number of collisions between enzyme and substrate.

The simplest model for the effect of condensates on enzymes is pure mass action: The reaction is enhanced when enzyme and substrate are co-concentrated in the condensate, but no changes are observed in the rate constant of the reaction. Accordingly, dramatic boosts have been observed in some enzyme-condensate systems. For example, doubling the concentration of the kinase FAK above its C_sat_ led to a 25-fold increase in the total auto-phosphorylation rate, which was attributed to an approximately 90.000-fold higher volume-normalized phosphorylation rate.^12^ Recruitment of the kinase MAPK3 to a SUMO:SIM condensate led to enhancement of phosphorylation rates up to ∼2-fold.^21^ These studies confirm that it is possible to enhance phosphorylation reactions through phase separation. However, the boost is rarely as large as predicted from mass action, indicating that the condensate environment is inhibitory in some way, which offsets the boost from co-partitioning.^45^ It is thus not surprising that we observe inhibition in the dense phase and this is therefore likely to be a common feature of condensates.

Inhibition could arise from changes in the enzymes as the hydrophobic condensate environment can lead to transient unfolding in condensates.^31^ Such effects are not necessarily inhibitory, but can also activate the enzyme, as observed for a lipase whose substrate-binding site becomes more accessible in the condensate or in organic co-solvents.^7^ In our case, the structure of the enzyme is practically unchanged in the condensate, ruling out structural changes as a source of inhibition. As the substrate is disordered, structural changes on the substrate side are also unlikely. These results thus indicate that the rate reduction originates in the emergent environment rather than in molecular changes.

Slow diffusion in condensates has been suggested as a generally inhibitory feature, although this has not been tested directly due to the difficulty in varying diffusion coefficients. Condensates form percolated networks that impose unique probe size dependence on dissociation. Proteins fall within the same size range as the pores in this meshwork, and the effective viscosity of the condensates varies with protein size.^38,39^ A unique aspect of the present study is the variation in the size of the molecule in which the protein is embedded. This allowed us to vary diffusion coefficient ∼50-fold without changing the rate constants of the enzyme:substrate interaction and uncovered a strong correlation between diffusion coefficient and the specificity constant of the phosphorylation reaction – providing a direct confirmation of the connection between slow diffusion and reduced enzyme catalysis. We were only successful in modulating diffusion rates up to a point, as substrates beyond a certain length all had similar diffusion rates. This was previously observed for IDPs of varied composition. Repeat proteins of different lengths have similar mean interaction strengths with the condensate scaffold, representing mainly the effect of chain length. This observation further supports the conclusion that disordered protein clients beyond a certain length will have similar diffusion coefficients in condensates.^40^

The longer substrates also partitioned more strongly into the condensates, as expected from the increased interactions between scaffold and client. This is not a problem in terms of the extraction of the rate constants, as K_P,S_ can be measured independently and accounted for in the modelling. However, it reveals an underlying limitation in ability to enhance enzymatic reactions through co-concentration in a condensate. Strong recruitment of a substrate requires favorable interaction between the substrate and the condensate scaffold. However, the strength of the interaction between client and scaffold controls the translational diffusion of the client.^23^ The expected boost from increased substrate partitioning into an enzyme-containing condensate is therefore tempered by the accompanying reduction in diffusion. This is likely to represent a general tradeoff in both natural and engineered enzyme-condensate reaction crucibles.

In conclusion, we have demonstrated a strong correlation between diffusion coefficients of substrates and enzyme kinetics within a condensate, which we attribute to mass transport limitations. Given the generally slow diffusion of macromolecules in the percolated network of a condensate, we suggest that such effects are likely to be common – especially for enzymes with macromolecular substrates. The correlation between diffusion and enzyme activity provides a direct link whereby the material properties of condensates affect the biochemical reactions that take place within them.

## Data availability

All data supporting the results of this study can be found in the article, supplementary, and source data files. Source Data are provided with this paper.

## ACKNOWLEDGEMENTS

This work was supported by grants from the Novo Nordisk Foundation (NNF20OC0063808, NNF23OC0082071), the Danish National Research Foundation (DNRF133, DNRF190). The “Biophysics and Biochemistry” and “Bioimaging” core facilities at Department of Molecular Biology & Genetics are thanked for technical support and instrument access. We acknowledge use of the Aarhus single molecule fluorescence (ASiMoF) infrastructure, funded by the Novo Nordisk Foundation (NNF20OC0061417). Authors would like to thank Mette Hoffmann Asmussen and Lisbeth S. Laursen for excellent technical support, and Paolo Arosio and Leila Margot Henkes for helpful comments on the manuscript.

