## Supplemental information for "Slow diffusion limits phosphorylation in a biomolecular condensate"

This file contains supplementary text describing the kinetic model used for fitting and numerical simulations in addition to 7 supplementary tables and 9 supplementary figures.

**Index:**

### Supplementary text:

Derivations of a rate law for a two-phase enzymatic system as a function of volume fraction of dense phase.

### Supplementary tables:

**Table 1:** Protein and peptide sequences used.

**Table 2:** Volume fractions of dense phase and reaction rate values obtained per substrate per condition.

**Table 3:** Partition coefficient values of labelled substrates and enzyme determined by confocal microscopy.

**Table 4:** Specificity constant values for each substrate in the dense and dilute phase derived from fitting.

**Table 5:** Fitted parameters from FRAP experiments of labelled substrates in [$Q_{5,8}$]-20 scaffold.

**Table 6:** Reaction rate values for mutant Substrate R-3K.

**Table 7:** Reaction rate values for increasing Enzyme concentrations.

### Supplementary figures:

**Figure 1:** Protein purity assessed by SDS-PAGE.

**Figure 2:** Temperature-dependent turbidity to evaluate the effect of substrates on phase behavior.

**Figure 3:** Phosphorylation of [$Q_{5,8}$]-24 $S_{WT}$ assessed by Phos-tag gel.

**Figure 4:** Hydrogen-deuterium exchange mass spectrometry data for MBD2-PKAc.

**Figure 5:** Time course of $^{32}$P-phosphorylated [$Q_{5,8}$]-x $S_{WT}$ substrates.

**Figure 6:** Decomposition of the contributions of phosphorylation reaction rates.

**Figure 7:** FRAP curves of 1 μM CF660R-[$Q_{5,8}$]-x $S_{WT}$ substrates recruited as client.

**Figure 8:** Time course of $^{32}$P-phosphorylated [$Q_{5,8}$]-24 $S_{R\text{-}3K}$ substrate.

**Figure 9:** Time course of phosphorylation at increasing enzyme concentrations.

### References

**SUPPLEMENTARY TEXT**

**Kinetic Model:**

**I.** Derivation of kinetic parameters for our model:

To assess the magnitude of the change introduced by condensates in the reaction rates, we calculate the ratio between the initial total rate in the biphasic/heterogenous system ($v_{Het}$) and in the homogeneous system ($v_{Hom}$), at the same total Substrate and Enzyme concentrations, and we call it rate enhancement (ξ):

$$\xi = \frac{v_{Het}}{v_{Hom}} \qquad (1)$$

We assume that in both systems the reaction scheme for our catalytical model is:

$$E + S \underset{k_{-1}}{\overset{k_1}{\longleftrightarrow}} ES \overset{k_2}{\rightarrow} EP \overset{k_3}{\rightarrow} E + P \qquad (2)$$

Where E is the enzyme, S the substrate, ES the intermediate enzyme-substrate complex, EP the intermediate enzyme-product complex, and P the product. The first step is reversible, $k_1$ is the association rate constant and $k_{-1}$ is the dissociation rate constant. The second and third steps are irreversible and correspond to phosphorylation ($k_2$ is the rate of phosphor-transfer) and product release ($k_3$ is the rate of ADP-release).

**a. An expression for $v_{Hom}$:**

If in the homogeneous system the reaction follows Michaelis-Menten kinetics, the rate of product formation follows:

$$v_{Hom} = \frac{k_{cat}[S][E]_0}{K_M + [S]} \qquad (3)$$

To derive a complete expression for $v_{Hom}$, we first consider Mass Action Law and obtain the following non-linear reaction equations:

$$\frac{d[S]}{dt} = -k_1[S][E] + k_{-1}[ES] \qquad (4)$$

$$\frac{d[E]}{dt} = -k_1[S][E] + k_{-1}[ES] + k_3[EP] \qquad (5)$$

$$\frac{d[ES]}{dt} = k_1[S][E] - k_{-1}[ES] - k_2[ES] \qquad (6)$$

$$\frac{d[EP]}{dt} = k_2[ES] - k_3[EP] \qquad (7)$$

$$\frac{d[P]}{dt} = k_3[EP] \qquad (8)$$

We consider the total enzyme concentration as constant (enzyme conservation):

$$[E]_0 = [E] + [ES] + [EP] = 0 \qquad (9)$$

And quasi-steady-state approximation for intermediate states:

$$\frac{d[ES]}{dt} = 0 \qquad (10)$$

$$\frac{d[EP]}{dt} = 0 \qquad (11)$$

Then:

$$\frac{d[ES]}{dt} = k_1[S][E] - k_{-1}[ES] - k_2[ES] = 0 \qquad (12)$$

$$k_1[S][E] = k_{-1}[ES] + k_2[ES] \qquad (13)$$

$$k_1[S][E] = (k_{-1} + k_2)[ES] \qquad (14)$$

$$[ES] = \frac{k_1[S][E]}{k_{-1} + k_2} \quad (15)$$

And:

$$\frac{d[EP]}{dt} = k_2[ES] - k_3[EP] = 0 \quad (16)$$

$$k_2[ES] = k_3[EP] \quad (17)$$

$$[EP] = \frac{k_2}{k_3}[ES] \quad (18)$$

Replacing [EP] in $[E]_0$:

$$[E]_0 = [E] + [ES] + [EP] = [E] + [ES] + \frac{k_2}{k_3}[ES] \quad (19)$$

$$[E]_0 = [E] + \left(1 + \frac{k_2}{k_3}\right)[ES] \quad (20)$$

So, solving [E]:

$$[E] = [E]_0 - \left(1 + \frac{k_2}{k_3}\right)[ES] \quad (21)$$

And replacing [E] in [ES] and rearranging:

$$[ES] = \frac{k_1\left([E]_0 - \left(1 + \frac{k_2}{k_3}\right)[ES]\right)[S]}{k_{-1} + k_2} \quad (22)$$

$$[ES](k_{-1} + k_2) = k_1[S]\left([E]_0 - \left(1 + \frac{k_2}{k_3}\right)[ES]\right) \quad (23)$$

$$[ES](k_{-1} + k_2) = k_1[S][E]_0 - k_1[S]\left(1 + \frac{k_2}{k_3}\right)[ES] \quad (24)$$

$$[ES](k_{-1} + k_2) + k_1[S]\left(1 + \frac{k_2}{k_3}\right)[ES] = k_1[S][E]_0 \quad (25)$$

$$[ES]\left(k_{-1} + k_2 + k_1[S]\left(1 + \frac{k_2}{k_3}\right)\right) = k_1[S][E]_0 \quad (26)$$

$$[ES] = \frac{k_1[S][E]_0}{k_{-1} + k_2 + k_1[S]\left(1 + \frac{k_2}{k_3}\right)} \quad (27)$$

Now, if

$$v_{Hom} = \frac{d[P]}{dt} \quad (28)$$

Then,

$$v_{Hom} = \frac{d[P]}{dt} = k_3[EP] = k_3\frac{k_2}{k_3}[ES] = \frac{k_2 k_1[S][E]_0}{k_{-1} + k_2 + k_1[S]\left(1 + \frac{k_2}{k_3}\right)} \quad (29)$$

If we multiply and divide by $k_1$, then by $k_3/(k_2 + k_3)$ and distribute, we get the expression for $v_{Hom}$:

$$v_{Hom} = \frac{\frac{k_2 k_3}{k_2 + k_3}[S][E]_0}{\left(\frac{k_3}{k_2 + k_3}\right)\left(\frac{k_{-1} + k_2}{k_1}\right) + [S]} \quad (30)$$

And our enzyme kinetic parameters take the form:

$$k_{cat} = \frac{k_2 k_3}{k_2 + k_3} \quad (31)$$

$$K_M = \left(\frac{k_3}{k_2 + k_3}\right)\left(\frac{k_{-1} + k_2}{k_1}\right) \quad (32)$$

$$K_{SP} = \frac{k_{cat}}{K_M} = \frac{\frac{k_2 k_3}{k_2 + k_3}}{\left(\frac{k_3}{k_2 + k_3}\right)\left(\frac{k_{-1} + k_2}{k_1}\right)} = \frac{k_2 k_1}{k_{-1} + k_2} \quad (33)$$

Where $K_{SP}$ corresponds to the specificity constant or catalytic efficiency of the enzyme, and we define its reciprocal of as:

$$R_K = \frac{1}{K_{SP}} \qquad (34)$$

For greater accuracy while fitting parameters in our simulations based on experimental data, we express $K_M$ as a function of $K_{SP}$ and $k_{cat}$ ([1]), therefore $v_{Hom}$ takes the final form:

$$v_{Hom} = \frac{C_{E,T}\ C_{S,T}}{R_{K,Hom} + \frac{C_{S,T}}{k_{cat,Hom}}} \qquad (35)$$

#### b. An expression for $v_{Het}$:

Now, we consider that the reaction takes place in a biphasic/heterogenous system. The total reaction rate adds the contributions of each phase weighed by their volume fraction (Φ), as follows:

$$v_{Het} = v_D\ \phi_D + v_B\ (1 - \phi_D) \quad (36)$$

Where the total volume fraction is:

$$\phi_T = \frac{V_D}{V_T} + \frac{V_B}{V_T} = \phi_D + \phi_B = 1 \quad (37)$$

Here, $V_D$ is the volume in the dense phase, $V_{dil}$ is the volume of the dilute phase and $V_T$ is the total volume of the system, so $\Phi_D$ is the volume fraction of the dense phase and $\Phi_{dil}$ is the volume fraction of the dilute phase.

According to Michaelis-Menten kinetics, the reaction rate in the dense phase takes this form:

$$v_D = \frac{k_{cat,D}\ C_{E,D}\ C_{S,D}}{K_{M,D} + C_{S,D}} \qquad (38)$$

And the reaction rate in the dilute phase is:

$$v_{dil} = \frac{k_{cat,dil}\ C_{E,dil}\ C_{S,dil}}{K_{M,dil} + C_{S,dil}} \quad (39)$$

These two reaction rates (38) and (39) depend on four unknown concentrations: Enzyme and Substrate concentrations in the dense and dilute phases, but they can be expressed in terms of: Total known concentrations ($C_T$), measurable partitioning coefficients ($K_P$) and volume fraction of the dense phase -which can also be measured and we consider a free variable in our analyses-. Below the analysis to get these final expressions for $v_D$ and $v_{dil}$.

If the principle of mass conservation applies in the heterogenous system, in the equilibrium, we have:

$$C_T V_T = C_D V_D + C_{dil} V_{dil} \qquad (40)$$

Dividing 40 by $V_T$:

$$C_T = C_D \phi_D + C_{dil} \phi_{dil} \quad (41)$$

And rearranging:

$$C_T - C_{dil}\phi_{dil} = C_D \phi_D \quad (42)$$

$$\frac{C_T - C_{dil}\phi_{dil}}{C_D} = \phi_D \qquad (43)$$

$$\frac{C_T}{C_D} - \frac{C_{dil}\phi_{dil}}{C_D} = \phi_D \qquad (44)$$

And replacing $\Phi_B$ according to (37):

$$\frac{C_T}{C_D} - \frac{dil}{C_D}(1 - \phi_D) = \phi_D \, (45)$$

$$\frac{C_T}{C_D} - \frac{C_{dil}}{C_D} + \frac{C_{dil}}{C_D}\phi_D = \phi_D \qquad (46)$$

$$\frac{C_T}{C_D} - \frac{C_{dil}}{C_D} = \phi_D - \frac{C_{dil}}{C_D}\phi_D \qquad (47)$$

$$\frac{C_T}{C_D} - \frac{C_{dil}}{C_D} = (1 - \frac{C_{dil}}{C_D})\phi_D \qquad (48)$$

$$\frac{C_T - C_{dil}}{C_D} = \frac{C_D - C_{dil}}{C_D}\phi_D \qquad (49)$$

We get:

$$\frac{C_T - C_{dil}}{C_D - C_{dil}} = \phi_D \qquad (50)$$

For a system where Substrate and Enzyme partition into both phases, we have that the partition coefficient of each component is:

$$K_P = \frac{C_D}{C_{dil}} \qquad (51)$$

If we reformulate (40) to express $C_D$ as a function of $K_P$ and $C_T$, then:

$$C_D = \frac{V_T}{V_D}C_T - \frac{V_{dil}}{V_D}C_{dil} \qquad (52)$$

Replacing $C_{dil}$ for (51):

$$C_D = \frac{V_T}{V_D}C_T - \frac{V_{dil}}{V_D}\frac{C_D}{K_P} \qquad (53)$$

$$C_D + \frac{V_{dil}}{V_D}\frac{C_D}{K_P} = \frac{V_T}{V_D}C_T \qquad (54)$$

$$C_D(1 + \frac{V_{dil}}{V_D\,K_P}) = \frac{V_T}{V_D}C_T \qquad (55)$$

$$C_D = \frac{\frac{V_T}{V_D}C_T}{1 + \frac{V_{dil}}{V_D\,K_P}} \qquad (56)$$

$$C_D = \frac{\frac{V_T}{V_D}C_T}{1 + \frac{V_{dil}}{V_D\,K_P}}\,\frac{K_P}{K_P} \qquad (57)$$

$$C_D = \frac{K_P\frac{V_T}{V_D}C_T}{K_P + \frac{V_{dil}}{V_D}} \qquad (58)$$

$$C_D = \frac{K_P\,V_T\,C_T}{K_P\,V_D + V_{dil}} \qquad (59)$$

And now, to express $C_B$ as a function of $K_P$ and $C_T$, we substitute $C_D$ in (58) with (51) and rearrange:

$$K_P\,C_{dil} = \frac{K_P\,V_T\,C_T}{K_P\,V_D + V_{dil}} \qquad (60)$$

$$C_{dil} = \frac{V_T\,C_T}{K_P\,V_D + V_{dil}} \qquad (61)$$

To get the final expressions for $v_D$, then we replace $C_D$ in (38) with (59):

$$v_D = \frac{k_{cat,D}\frac{K_{P,E}\,V_T\,C_{E,T}}{K_{P,E}\,V_D + V_{dil}}\,\frac{K_{P,S}\,V_T\,C_{S,T}}{K_{P,S}\,V_D + V_{dil}}}{K_{M,D} + \frac{K_{P,S}\,V_T\,C_{S,T}}{K_{P,S}\,V_D + V_{dil}}} \qquad (62)$$

$$v_D = \frac{k_{cat,D}\,{V_T}^2\,\frac{K_{P,E}\,C_{E,T}}{K_{P,E}\,V_D + V_{dil}}\,\frac{K_{P,S}\,C_{S,T}}{K_{P,S}\,V_D + V_{dil}}}{K_{M,D} + \frac{K_{P,S}\,V_T\,C_{S,T}}{K_{P,S}\,V_D + V_{dil}}} \qquad (63)$$

For $v_{dil}$, in the dilute phase, we replace $C_{dil}$ in (39) with (61):

$$v_{dil} = \frac{k_{cat,dil} \frac{V_T\, C_{E,T}}{K_{P,E}\, V_D + V_{dil}} \frac{V_T\, C_{S,T}}{K_{P,S}\, V_D + V_{dil}}}{K_{M,dil} + \frac{V_T\, C_{S,T}}{K_{P,S}\, V_D + V_{dil}}} \quad (64)$$

$$v_{dil} = \frac{k_{cat,dil}\ {V_T}^2 \frac{C_{E,T}}{K_{P,E}\, V_D + V_{dil}} \frac{C_{S,T}}{K_{P,S}\, V_D + V_{dil}}}{K_{M,dil} + \frac{V_T\, C_{S,T}}{K_{P,S}\, V_D + V_{dil}}} \quad (65)$$

The total reaction rate in the heterogeneous phase then follows:

$$v_{Het} = \frac{k_{cat,D}\ {V_T}^2 \frac{K_{P,E}\, C_{E,T}}{K_{P,E}\, V_D + V_{dil}} \frac{K_{P,S}\, C_{S,T}}{K_{P,S}\, V_D + V_{dil}}}{K_{M,D} + \frac{K_{P,S}\, V_T\, C_{S,T}}{K_{P,S}\, V_D + V_{dil}}}\ \phi_D + \frac{k_{cat,dil}\ {V_T}^2 \frac{C_{E,T}}{K_{P,E}\, V_D + V_{dil}} \frac{C_{S,T}}{K_{P,S} * V_D + V_{dil}}}{K_{M,B} + \frac{V_T\, C_{S,T}}{K_{P,S}\, V_D + V_{dil}}}\ (1 - \phi_D) \quad (66)$$

And if we divide and multiply by $k_{cat}$, then we have:

$$v_{Het} = \frac{{V_T}^2 \frac{K_{P,E}\, C_{E,T}}{K_{P,E}\, V_D + V_{dil}} \frac{K_{P,S}\, C_{S,T}}{K_{P,S}\, V_D + V_{dil}}}{R_{K,D} + \frac{\frac{K_{P,S}\, V_T\, C_{S,T}}{K_{P,S}\, V_D + V_{dil}}}{k_{cat,D}}}\ \phi_D + \frac{{V_T}^2 \frac{C_{E,T}}{K_{P,E}\, V_D + V_{dil}} \frac{C_{S,T}}{K_{P,S}\, V_D + V_{dil}}}{R_{K,dil} + \frac{\frac{V_T\, C_{S,T}}{K_{P,S}\, V_D + V_{dil}}}{k_{cat,dil}}}\ (1 - \phi_D) \quad (67)$$

### Supplementary Table 1:

| # | Short name | Peptide or Protein sequence | aa | Function | Mol Weight (Da) | ε ($M^{-1}cm^{-1}$) |
|---|---|---|---|---|---|---|
| 1 | Kemptide | LRRA**S**LGGY | 9 | Substrate | 992.12 | 1490 |
| 2 | $[Q_{5,8}]$-2 $S_{WT}$ | GRGDQPYQLRRA**S**LGGRGDQPYQ | 23 | Substrate | 2575.79 | 2980 |
| 3 | $[Q_{5,8}]$-4 $S_{WT}$ | GRGDQPYQGRGDQPYQLRRA**S**LGGRGDQPYQGRGDQPYQ | 39 | Substrate | 4379.65 | 5960 |
| 4 | $[Q_{5,8}]$-10 $S_{WT}$ | MGSSHHHHHHSSGLVPRGSHMSKGPGRGDQPYQGRGDQPYQG<br>RGDQPYQGRGDQPYQGRGDQPYQLRRA**S**LGGRGDQPYQGRGD<br>QPYQGRGDQPYQGRGDQPYQGRGDQPYQGY | 114 | Substrate | 12675.42 | 16390 |
| 5 | $[Q_{5,8}]$-14 $S_{WT}$ | MGSSHHHHHHSSGLVPRGSHMSKGPGRGDQPYQGRGDQPYQG<br>RGDQPYQGRGDQPYQGRGDQPYQGRGDQPYQGRGDQPYQLRR<br>A**S**LGGRGDQPYQGRGDQPYQGRGDQPYQGRGDQPYQGRGDQP<br>YQGRGDQPYQGRGDQPYQGY | 146 | Substrate | 16283.16 | 22350 |
| 6 | $[Q_{5,8}]$-20 $S_{WT}$ | MGSSHHHHHHSSGLVPRGSHMSKGPGRGDQPYQGRGDQPYQG<br>RGDQPYQGRGDQPYQGRGDQPYQGRGDQPYQGRGDQPYQGRG<br>DQPYQGRGDQPYQGRGDQPYQLRRA**S**LGGRGDQPYQGRGDQP<br>YQGRGDQPYQGRGDQPYQGRGDQPYQGRGDQPYQGRGDQPYQ<br>GRGDQPYQGRGDQPYQGRGDQPYQGY | 194 | Substrate | 21694.76 | 31290 |
| 7 | $[Q_{5,8}]$-24 $S_{WT}$ | MGSSHHHHHHSSGLVPRGSHMSKGPGRGDQPYQGRGDQPYQG<br>RGDQPYQGRGDQPYQGRGDQPYQGRGDQPYQGRGDQPYQGRG<br>DQPYQGRGDQPYQGRGDQPYQGRGDQPYQGRGDQPYQLRRA**S**<br>LGGRGDQPYQGRGDQPYQGRGDQPYQGRGDQPYQGRGDQPYQ<br>GRGDQPYQGRGDQPYQGRGDQPYQGRGDQPYQGRGDQPYQGR<br>GDQPYQGRGDQPYQGY | 226 | Substrate | 25302.50 | 37250 |
| 8 | $[Q_{5,8}]$-24 $S_{R-2K}$ | MGSSHHHHHHSSGLVPRGSHMSKGPGRGDQPYQGRGDQPYQG<br>RGDQPYQGRGDQPYQGRGDQPYQGRGDQPYQGRGDQPYQGRG<br>DQPYQGRGDQPYQGRGDQPYQGRGDQPYQGRGDQPYQLRKA**S**<br>LGGRGDQPYQGRGDQPYQGRGDQPYQGRGDQPYQGRGDQPYQ<br>GRGDQPYQGRGDQPYQGRGDQPYQGRGDQPYQGRGDQPYQGR<br>GDQPYQGRGDQPYQGY | 226 | Substrate | 25274.49 | 37250 |
| 9 | $[Q_{5,8}]$-24 $S_{R-3K}$ | MGSSHHHHHHSSGLVPRGSHMSKGPGRGDQPYQGRGDQPYQG<br>RGDQPYQGRGDQPYQGRGDQPYQGRGDQPYQGRGDQPYQGRG<br>DQPYQGRGDQPYQGRGDQPYQGRGDQPYQGRGDQPYQLKRA**S**<br>LGGRGDQPYQGRGDQPYQGRGDQPYQGRGDQPYQGRGDQPYQ<br>GRGDQPYQGRGDQPYQGRGDQPYQGRGDQPYQGRGDQPYQGR<br>GDQPYQGRGDQPYQGY | 226 | Substrate | 25274.49 | 37250 |
| 10 | $[Q_{5,8}]$-30 $S_{WT}$ | MGSSHHHHHHSSGLVPRGSHMSKGPGRGDQPYQGRGDQPYQG<br>RGDQPYQGRGDQPYQGRGDQPYQGRGDQPYQGRGDQPYQGRG<br>DQPYQGRGDQPYQGRGDQPYQGRGDQPYQGRGDQPYQGRGDQ<br>PYQGRGDQPYQGRGDQPYQLRRA**S**LGGRGDQPYQGRGDQPYQ<br>GRGDQPYQGRGDQPYQGRGDQPYQGRGDQPYQGRGDQPYQGR<br>GDQPYQGRGDQPYQGRGDQPYQGRGDQPYQGRGDQPYQGRGD<br>QPYQGRGDQPYQGRGDQPYQGY | 274 | Substrate | 30714.10 | 46190 |
| 11 | $[Q_{5,8}]$-40 $S_{WT}$ | MGSSHHHHHHSSGLVPRGSHMSKGPGRGDQPYQGRGDQPYQG<br>RGDQPYQGRGDQPYQGRGDQPYQGRGDQPYQGRGDQPYQGRG<br>DQPYQGRGDQPYQGRGDQPYQGRGDQPYQGRGDQPYQGRGDQ<br>PYQGRGDQPYQGRGDQPYQGRGDQPYQGRGDQPYQGRGDQPY<br>QGRGDQPYQGRGDQPYQLRRA**S**LGGRGDQPYQGRGDQPYQGR<br>GDQPYQGRGDQPYQGRGDQPYQGRGDQPYQGRGDQPYQGRGD<br>QPYQGRGDQPYQGRGDQPYQGRGDQPYQGRGDQPYQGRGDQP<br>YQGRGDQPYQGRGDQPYQGRGDQPYQGRGDQPYQGRGDQPYQ<br>GRGDQPYQGRGDQPYQGY | 354 | Substrate | 39733.44 | 61090 |
| 12 | $[Q_{5,8}]$-20 | MGSSHHHHHHSSGLVPRGSHMSKGPGRGDQPYQGRGDQPYQG<br>RGDQPYQGRGDQPYQGRGDQPYQGRGDQPYQGRGDQPYQGRG<br>DQPYQGRGDQPYQGRGDQPYQGRGDQPYQGRGDQPYQGRGDQ<br>PYQGRGDQPYQGRGDQPYQGRGDQPYQGRGDQPYQGRGDQPY<br>QGRGDQPYQGRGDQPYQGY | 187 | Scaffold | 20940.86 | 31290 |
| 13 | $[Q_{5,8}]$-20-p66α | MGSSHHHHHHSSGLVPRGSHMSKGPGRGDQPYQGRGDQPYQG<br>RGDQPYQGRGDQPYQGRGDQPYQGRGDQPYQGRGDQPYQGRG<br>DQPYQGRGDQPYQGRGDQPYQTSPEERERMIKQLKEELRLEE<br>AKLVLLKKLRQSQIQKEATAQKGRGDQPYQGRGDQPYQGRGD<br>QPYQGRGDQPYQGRGDQPYQGRGDQPYQGRGDQPYQGRGDQP<br>YQGRGDQPYQGRGDQPYQGY | 230 | Scaffold | 26071.89 | 31290 |
| 14 | MBD2-PKAc (PKA) | MGSSHHHHHHSSGLVPRGSHMVTDEDIRKQEER**V**QQVRKKLE<br>EALMADASQSGTGNAAAAKKGSEQESVKEFLAKAKEDFLKKW<br>ETPSQNTAQLDQFDRIKTLGTGSFGRVMLVKHKESGNHYAMK<br>ILDKQKVVKLKQIEHTLNEKRILQAVNFPFLVKLEFSFKDNS<br>NLYMVMEYVAGGEMFSHLRRIGRFSEPHARFYAAQIVLTFEY<br>LHSLDLIYRDLKPENLLIDQQGYIQVTDFGFAKRVKGRTWTL<br>CGTPEYLAPEIILSKGYNKAVDWWALGVLIYEMAAGYPPFFA<br>DQPIQIYEKIVSGKVRFPSHFSSDLKDLLRNLLQVDLTKRFG<br>NLKNGVNDIKNHKWFATTDWIAIYQRKVEAPFIPKFKGPGDT<br>SNFDDYEEEEIRVSINEKCGKEFTEF | 404 | Enzyme | 46505.12 | 53860 |

Color coding for **Supplementary Table 1**:

- 6xHis-tag
- Thrombin cleavage sequence
- Variable/extra sequence
- Octapeptide
- PKA substrate motif, catalytic serine is shown in **bold**
- p66α sequence
- MBD2 sequence

**Supplementary Table 2:** Volume fractions of dense phase and reaction rate values obtained per substrate per condition

| Substrate | Scaffold Concentration (μM) | $\Phi_D$ | Reaction rate Homogeneous $v_{Hom}$ (min$^{-1}$) | Reaction rate Heterogeneous $v_{Het, Total}$ (min$^{-1}$) | Reaction rate Heterogeneous $v_{Het, dil}$ (min$^{-1}$) |
|---|---|---|---|---|---|
| Kemptide | 0 | | 5.31 ± 4.07 | | |
| | 5 | 0.0001 | | 16.50 ± 8.36 | 17.27 ± 10.29 |
| | 10 | 0.0003 | | 19.04 ± 3.05 | 15.12 ± 5.17 |
| | 25 | 0.0021 | | 28.92 ± 16.29 | 16.81 ± 8.52 |
| | 50 | 0.0283 | | 30.22 ± 9.77 | 13.46 ± 8.10 |
| | 100 | 0.0669 | | 28.42 ± 8.03 | 12.44 ± 3.58 |
| [$Q_{5,8}$]-2 $S_{WT}$ | 0 | | 12.27 ± 6.54 | | |
| | 5 | 0.0001 | | 45.50 ± 12.42 | 18.29 ± 4.75 |
| | 10 | 0.0003 | | 49.03 ± 11.43 | 19.42 ± 8.73 |
| | 25 | 0.0021 | | 49.97 ± 8.88 | 12.10 ± 8.41 |
| | 50 | 0.0283 | | 51.15 ± 20.16 | 5.23 ± 2.58 |
| | 100 | 0.0669 | | 35.64 ± 8.86 | 3.11 ± 2.59 |
| [$Q_{5,8}$]-4 $S_{WT}$ | 0 | | 26.64 ± 9.91 | | |
| | 5 | 0.0001 | | 55.25 ± 11.80 | 32.86 ± 5.46 |
| | 10 | 0.0003 | | 56.30 ± 10.65 | 34.36 ± 9.93 |
| | 25 | 0.0021 | | 59.05 ± 11.40 | 18.40 ± 6.57 |
| | 50 | 0.0283 | | 49.97 ± 6.35 | 13.32 ± 4.49 |
| | 100 | 0.0669 | | 34.34 ± 9.14 | 13.83 ± 8.81 |
| [$Q_{5,8}$]-10 $S_{WT}$ | 0 | | 124.72 ± 20.16 | | |
| | 5 | 0.0001 | | 206.96 ± 20.80 | 125.88 ± 8.16 |
| | 10 | 0.0003 | | 224.04 ± 28.28 | 143.04 ± 8.80 |
| | 25 | 0.0021 | | 244.76 ± 27.00 | 127.92 ± 15.72 |
| | 50 | 0.0283 | | 209.32 ± 22.56 | 84.48 ± 14.20 |
| | 100 | 0.0669 | | 115.96 ± 13.20 | 50.40 ± 8.76 |
| [$Q_{5,8}$]-14 $S_{WT}$ | 0 | | 139.65 ± 30.15 | | |
| | 5 | 0.0001 | | 249.20 ± 22.70 | 86.40 ± 18.65 |
| | 10 | 0.0003 | | 257.50 ± 38.35 | 121.35 ± 44.90 |
| | 25 | 0.0021 | | 266.95 ± 51.60 | 75.70 ± 17.90 |
| | 50 | 0.0283 | | 205.70 ± 53.15 | 27.17 ± 2.33 |
| | 100 | 0.0669 | | 57.35 ± 13.55 | 15.93 ± 8.82 |
| [$Q_{5,8}$]-20 $S_{WT}$ | 0 | | 89.11 ± 16.56 | | |
| | 5 | 0.0001 | | 120.11 ± 19.50 | 59.78 ± 13.77 |
| | 10 | 0.0003 | | 118.94 ± 23.39 | 63.83 ± 16.93 |
| | 25 | 0.0021 | | 122.56 ± 21.17 | 30.94 ± 13.85 |
| | 50 | 0.0283 | | 101.00 ± 26.39 | 33.20 ± 3.02 |
| | 100 | 0.0669 | | 27.82 ± 9.59 | 1.91 ± 3.82 |
| [$Q_{5,8}$]-24 $S_{WT}$ | 0 | | 101.80 ± 24.70 | | |
| | 5 | 0.0001 | | 135.45 ± 26.20 | 14.49 ± 10.89 |
| | 10 | 0.0003 | | 144.10 ± 19.15 | 27.38 ± 12.44 |
| | 25 | 0.0021 | | 99.75 ± 35.70 | 7.71 ± 10.61 |
| | 50 | 0.0283 | | 69.60 ± 16.60 | 8.01 ± 10.38 |
| | 100 | 0.0669 | | 21.57 ± 12.44 | 2.57± 5.14 |
| [$Q_{5,8}$]-30 $S_{WT}$ | 0 | | 105.00 ± 15.00 | | |
| | 5 | 0.0001 | | 96.35 ± 13.00 | 10.96 ± 13.87 |
| | 10 | 0.0003 | | 103.30 ± 15.00 | 19.93 ± 15.78 |
| | 25 | 0.0021 | | 71.25 ± 17.70 | 8.77 ± 7.23 |
| | 50 | 0.0283 | | 62.15 ± 17.75 | 9.08 ± 2.81 |
| | 100 | 0.0669 | | 17.07 ± 4.48 | 1.60 ± 3.16 |
| [$Q_{5,8}$]-40 $S_{WT}$ | 0 | | 48.36 ± 10.88 | | |

Volume fraction of the dense phase for each [$Q_{5,8}$]-20 + [$Q_{5,8}$]-20-p66$\alpha$ (9:1) scaffold concentration used in our experiments. Phosphorylation reaction rate values were obtained from global fitting of two experimental determinations (n=2) per condition. The error bars correspond to the propagated standard error of the rates.

**Supplementary Table 3:** Partition coefficient values of labelled substrates and enzyme determined by fluorescence confocal microscopy.

| Clients | $K_P$ |
|---|---|
| Kemptide | 4.6 ± 0.7 |
| [$Q_{5,8}$]-2 $S_{WT}$ | 12.4 ± 2.2 |
| [$Q_{5,8}$]-4 $S_{WT}$ | 31.4 ± 3.6 |
| [$Q_{5,8}$]-10 $S_{WT}$ | 431 ± 48.2 |
| [$Q_{5,8}$]-14 $S_{WT}$ | 380 ± 90.1 |
| [$Q_{5,8}$]-20 $S_{WT}$ | 764 ± 205 |
| [$Q_{5,8}$]-24 $S_{WT}$ | 1503 ± 110 |
| [$Q_{5,8}$]-30 $S_{WT}$ | 1590 ± 245 |
| MBD2-PKAc | 46.6 ± 5.2 |

Partition coefficient values of client molecules in [$Q_{5,8}$]-20 + [$Q_{5,8}$]-20-p66α (9:1) scaffold. $K_P$ values are the average of ten individual point measurements per client molecule (n=10). Errors correspond to the standard error of the mean. The numbers are reported with high digital precision on purpose.

**Supplementary Table 4:** Specificity constant values for each substrate in the dense and dilute phase derived from fitting.

| Clients | $K_{SP,D}$ ($\mu M^{-1} s^{-1}$) | $K_{SP,dil}$ ($\mu M^{-1} s^{-1}$) |
|---|---|---|
| Kemptide | 7.37 ± 2.03 | 5.24 ± 0.74 |
| $[Q_{5,8}]$-2 $S_{WT}$ | 1.49 ± 0.31 | 5.53 ± 0.51 |
| $[Q_{5,8}]$-4 $S_{WT}$ | 0.29 ± 0.11 | 3.04 ± 0.82 |
| $[Q_{5,8}]$-10 $S_{WT}$ | 0.147 ± 0.023 | 3.69 ± 0.68 |
| $[Q_{5,8}]$-14 $S_{WT}$ | 0.069 ± 0.017 | 3.86 ± 0.86 |
| $[Q_{5,8}]$-20 $S_{WT}$ | 0.057 ± 0.014 | 3.61 ± 1.00 |
| $[Q_{5,8}]$-24 $S_{WT}$ | 0.040 ± 0.015 | 7.35 ± 0.61 |
| $[Q_{5,8}]$-30 $S_{WT}$ | 0.026 ± 0.006 | 3.42 ± 1.03 |
| $[Q_{5,8}]$-24 $S_{R\text{-}3K}$ | 0.001 ± 0.001 | 0.022 ± 0.006 |

Specificity constants were obtained from fitting to our model with partitioning coefficients constrained to values obtained experimentally. $K_{SP,Hom}$ for WT and R-3K Substrates was obtained from ([6]). The numbers are reported with high digital precision on purpose.

**Supplementary Table 5:** Fitted parameters from FRAP experiments of labelled substrates in $[Q_{5,8}]$-20 scaffold.

| Substrate | Normalized $F_{mobile}$ | τ (sec) | $D_{i,app}$ [a] |
|---|---|---|---|
| Kemptide | 1.01 ± 0.03 | 1.23 ± 0.02 | 1.60 ± 0.027 |
| $[Q_{5,8}]$-2 $S_{WT}$ | 1.03 ± 0.01 | 5.27 ± 0.01 | 0.372 ± 0.001 |
| $[Q_{5,8}]$-4 $S_{WT}$ | 0.97 ± 0.01 | 6.88 ± 0.07 | 0.285 ± 0.003 |
| $[Q_{5,8}]$-10 $S_{WT}$ | 1.09 ± 0.02 | 23.6 ± 0.40 | 0.072 ± 0.001 |
| $[Q_{5,8}]$-14 $S_{WT}$ | 0.99 ± 0.03 | 28.4 ± 0.27 | 0.069 ± 0.001 |
| $[Q_{5,8}]$-20 $S_{WT}$ | 0.93 ± 0.06 | 25.3 ± 0.41 | 0.078 ± 0.001 |
| $[Q_{5,8}]$-24 $S_{WT}$ | 0.76 ± 0.04 | 19.9 ± 0.27 | 0.099 ± 0.001 |
| $[Q_{5,8}]$-30 $S_{WT}$ | 0.78 ± 0.05 | 27.4 ± 0.20 | 0.072 ± 0.001 |

Values corresponding to normalized mobile fraction, recovery time and apparent diffusion coefficient values derived from fitting FRAP curves (n=3) using single exponential model. Error correspond to the standard error of the mean.

**Supplementary Table 6:** Reaction rate values for mutant Substrate R-3K.

| Substrate | Scaffold Concentration (μM) | $\Phi_D$ | Reaction rate Homogeneous $v_{Hom}$ ($min^{-1}$) | Reaction rate Heterogeneous $v_{Het, Total}$ ($min^{-1}$) |
|---|---|---|---|---|
| $[Q_{5,8}]$-24 $S_{R-3K}$ | 0 | | 3.39 ± 0.41 | |
| | | | 3.09 ± 0.39 | |
| | 5 | 0.0001 | | 6.02 ± 1.29 |
| | 10 | 0.0003 | | 6.80 ± 0.71 |
| | 25 | 0.0021 | | 6.27 ± 0.83 |
| | 50 | 0.0283 | | 5.64 ± 0.74 |
| | 100 | 0.0669 | | 3.26 ± 0.69 |

Phosphorylation reaction rate values were obtained from global fitting of two experimental determinations (n=2) per condition. For the homogeneous system the reaction rate was measured in duplicate two times (n=4). The error bars correspond to the propagated standard error of the rates.

**Supplementary Table 7:** Reaction rates for increasing enzyme concentrations.

| Substrate | Enzyme Concentration (nM) | Reaction rate Heterogeneous $v_{Het, Total}$ ($min^{-1}$) |
|---|---|---|
| [$Q_{5,8}$]-2 $S_{WT}$ | 0.2 | 61.5 ± 12.60 |
| | 0.4 | 46.0 ± 9.15 |
| | 0.8 | 40.2 ± 8.21 |
| | 1.2 | 40.6 ± 4.43 |
| [$Q_{5,8}$]-20 $S_{WT}$ | 0.2 | 79.4 ± 14.88 |
| | 0.4 | 56.2 ± 6.35 |
| | 0.8 | 39.0 ± 8.50 |
| | 1.2 | 36.4 ± 6.14 |

Phosphorylation reaction rate values were obtained from global fitting of two experimental determinations (n=2) per condition. The error bars correspond to the propagated standard error of the rates.

**Supplementary Figure 1:**

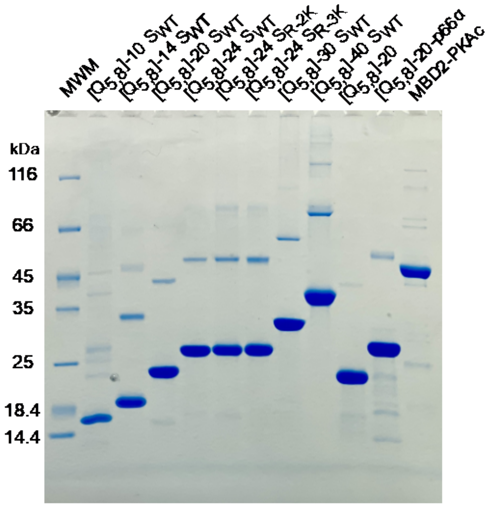


**Protein purity assessed by SDS-PAGE.** Gradient gel of all the purified recombinant proteins used in this work (purity >90%). For each lane 30 µg of protein were loaded, except for MDB2-PKAc were only 15 µg was loaded. For substrates and scaffold proteins there is second band with apparent double molecular weight, which corresponds to unspecific interactions of these proteins with SDS in the gel.

**Supplementary Figure 2:**

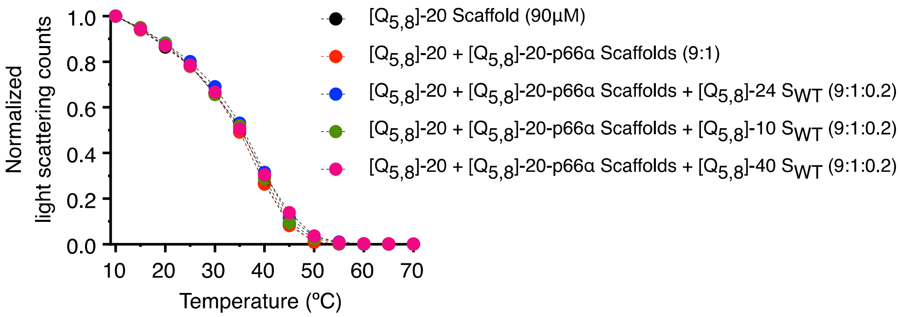


**Temperature-dependent turbidity experiments to evaluate the effect of substrates on phase behavior.** [$Q_{5,8}$]-20 scaffold was doped with [$Q_{5,8}$]-20-p66α scaffold at a molar ratio of 9:1, and then with [$Q_{5,8}$]-10$S_{WT}$, [$Q_{5,8}$]-24$S_{WT}$ and [$Q_{5,8}$]-40$S_{WT}$. The addition of [$Q_{5,8}$]-20-p66α as well as the substrates at selected concentrations showed no significant influence on the phase behavior of the [$Q_{5,8}$]-20 scaffold.

**Supplementary Figure 3:**

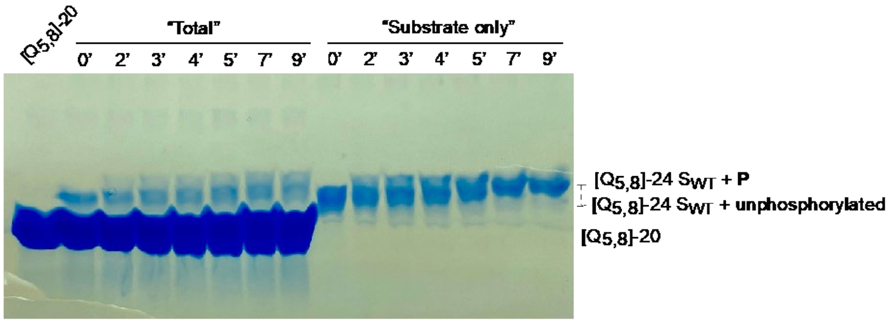


**Phosphorylation of [$Q_{5,8}$]-24 $S_{WT}$ assessed by Phos-tag gel.** Phosphorylation reaction of substrate [$Q_{5,8}$]-24 $S_{WT}$ in presence and absence of scaffold [$Q_{5,8}$]-20, at different time points. Equivalent amounts of substrate were loaded in each well. In presence if scaffold ("Total") phosphorylation is incomplete after 9 minutes, showing a slurry band corresponding to different degree of phosphorylation of the substrate. Phosphorylation in absence of scaffold ("Substrate only") shows complete reaction after 9 minutes, evidenced by the gap difference of the full protein substrate band between initial (0') and final (9') time steps.

**Supplementary Figure 4:**

**A)**

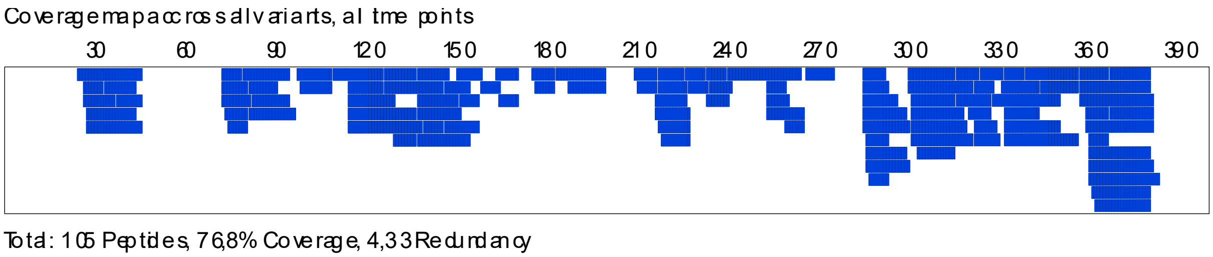


**B)**

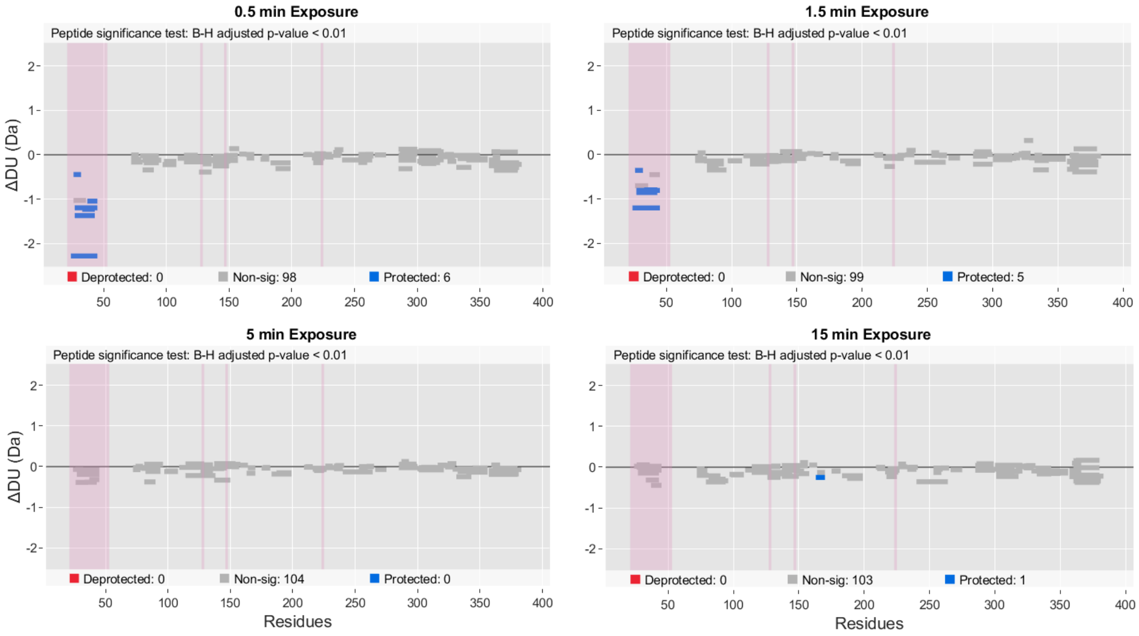


**Hydrogen-deuterium exchange mass spectrometry data for MBD2-PKAc**. **A)** Coverage map of MBD2-PKAc, overlayed from all variants using DynamX. Mass spectrometry analyzed 105 peptides which covered 76.8% of 404 amino acids in the sequence, with 4.33 in average of redundancy. **B)** Significancy test of deuterium uptake between MBD2-PKA with and without condensates with 99% confidential using Deuteros. The software found 6 and 5 peptides significantly more protected in the 0.5- and 1.5-min exposure time, located at the beginning of the construct, which belong to MBD2 (highlighted in pink). Regions in the middle of the sequence that are also highlighted belong to the residues in the catalytic site, which did not show a significant uptake difference.

**Supplementary Figure 5:**

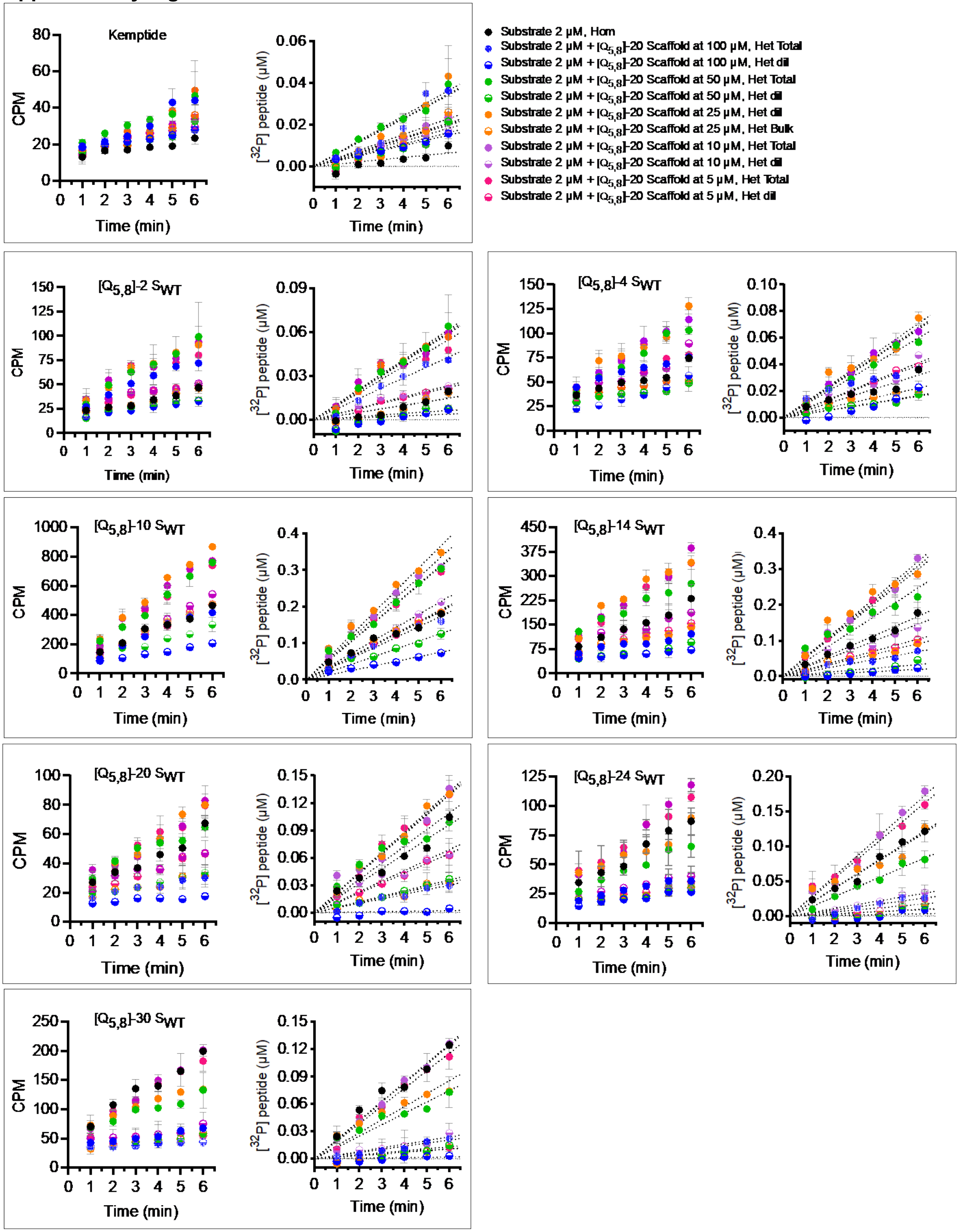


**Time course of $^{32}$P-phosphorylated [$Q_{5,8}$]-x $S_{WT}$ substrates** (at 2 μM) in homogeneous solution and a biphasic system (Het, Total) and dilute phase (Het, dil) at increasing scaffold concentrations. Each measurement was done by duplicate (n=2) and error bars correspond to the standard deviation.

**Supplementary Figure 6:**

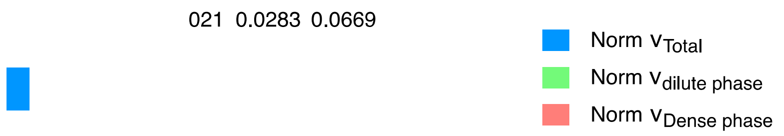


**Decomposition of the contributions of phosphorylation reaction rates** in the dense ($v_{Het,D}$ ) and dilute phase ($v_{Het,\ dil}$) to the total ($v_{Het,Total}$) normalized to 1, for all the substrates at different $\Phi_D$. For the longest substrates, the contribution of the dilute phase is negligible, and the dense phase is dominant. The contribution of the dilute phase increases with for short substrates (weaker partitioning). The height of the bar represents the normalized (to the total) in vitro phosphorylation reaction rate for each condition, and the error bars correspond to the propagated standard error of the rates.

**Supplementary Figure 7:**

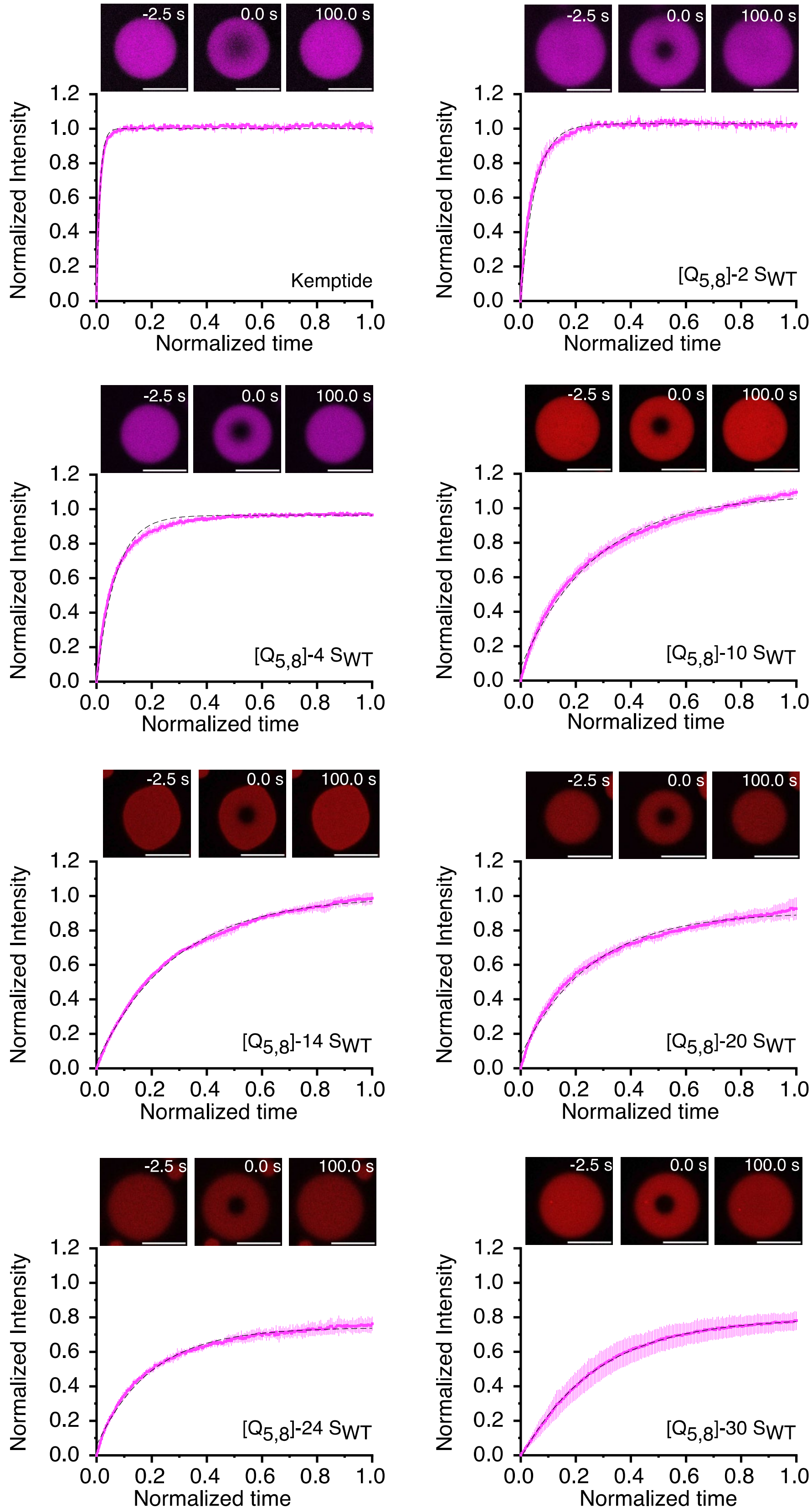


**FRAP curves of 1 μM CF660R-[$Q_{5,8}$]-x $S_{WT}$ substrates recruited as client** into 50 μM [$Q_{5,8}$]-20 scaffold (scale bars correspond to 10 μm). Three independent FRAP-determinations (n=3) were performed per substrate and averaged. Dashed line represents the fit to a simple exponential recovery model.

**Supplementary Figure 8:**

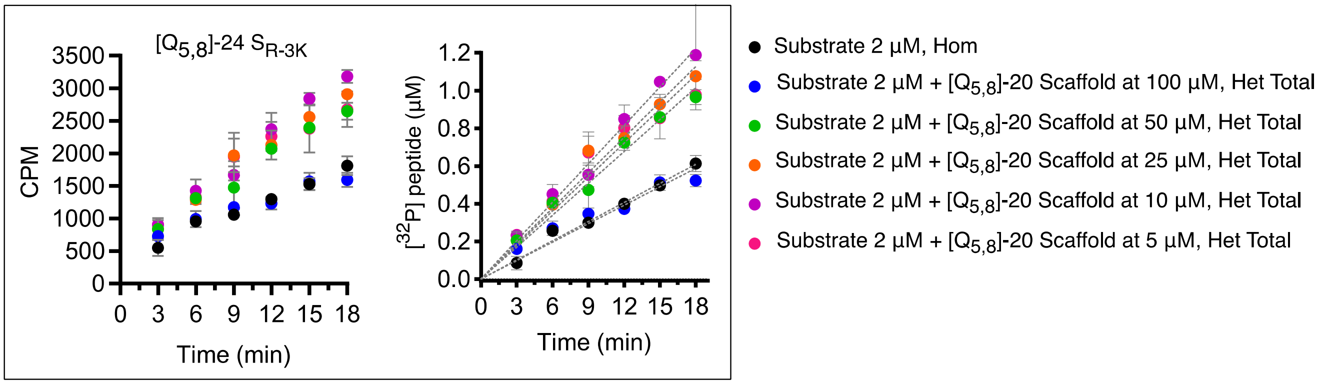


**Time course of $^{32}$P-phosphorylated [$Q_{5,8}$]-24 $S_{R\text{-}3K}$ substrate** (at 2 µM) in homogeneous solution and a biphasic system (Het, Total) and dilute phase (Het, dil) at increasing scaffold concentrations. Each measurement was done by duplicate (n=2) and error bars correspond to the standard deviation.

**Supplementary Figure 9:**

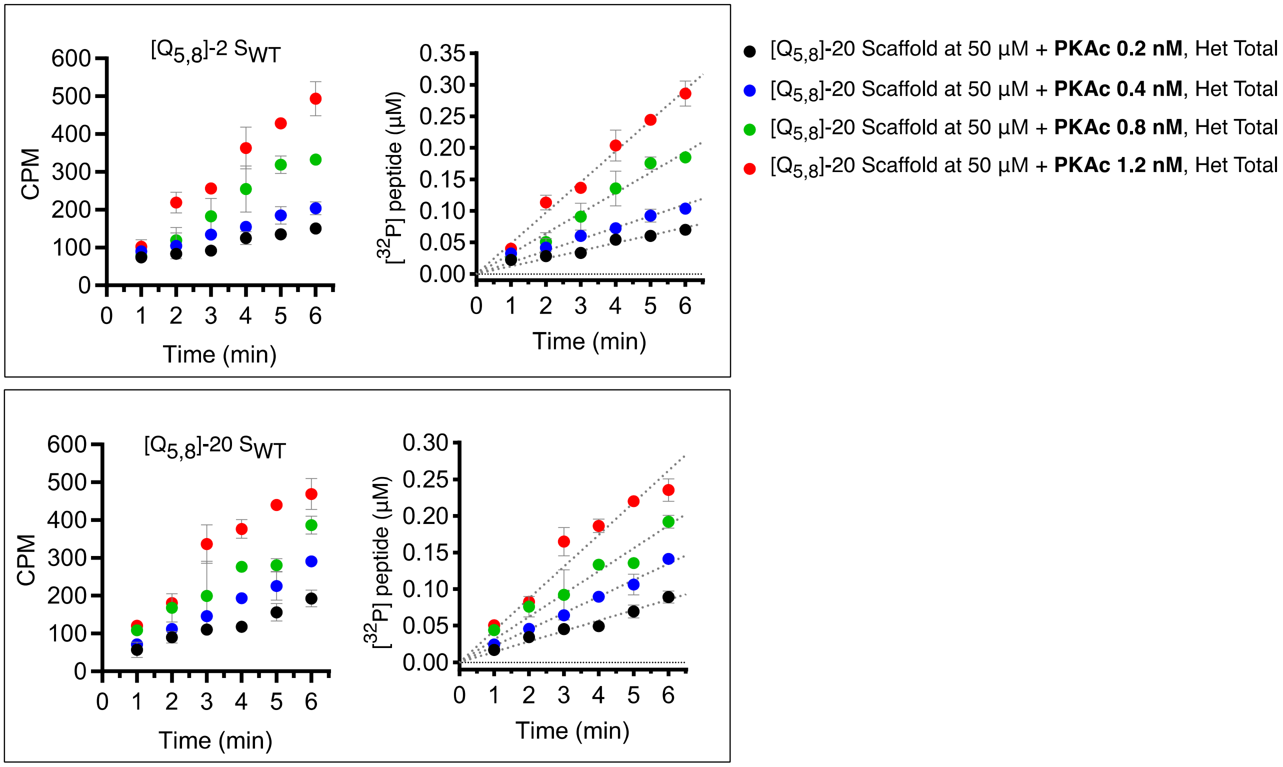


**Time course of $^{32}$P-phosphorylated [$Q_{5,8}$]-2 $S_{WT}$ and [$Q_{5,8}$]-20 $S_{WT}$ substrates at increasing enzyme concentrations.** Substrate concentration at 2 μM in homogeneous solution and a biphasic system (Het, Total) and dilute phase (Het, dil) at 50 μM scaffold concentration. Each measurement was done by duplicate (n=2) and error bars correspond to the standard deviation.
